# From Peaks to Power: Systematic Evaluation of Chromatographic Sampling Reveals Determinants of Quantification and Biological Discovery in DIA Proteomics

**DOI:** 10.64898/2026.05.13.724964

**Authors:** Lee S. Cantrell, Seth Just, Alexey Stukalov, Omid C. Farokhzad, Serafim Batzoglou

**Affiliations:** Seer, Inc, Redwood City, CA, USA

**Keywords:** Quantitative Proteomics, Accuracy, Datapoints Per Peak, Data-Independent Acquisition, Statistical Power, Population Proteomics, Biological Resolution

## Abstract

Modern DIA proteomics increasingly emphasizes throughput and depth for large-cohort studies, but methods are often optimized using proxy metrics that can mask losses in quantifiable signal and statistical power. Here, we evaluate how datapoints per peak and other chromatographic features jointly contribute to quantification and downstream biological discovery. Using a matrix-matched calibration curve dataset, we checked how the number of datapoints per peak (DPPP) affects the limits of detection and quantification (LOD/LOQ). Reduced DPPP minimally affected LOD but substantially degraded LOQ. Feature modeling and nonparametric association analyses identified precursor peak area as the strongest feature-level predictor of LOQ, whereas DPPP showed weaker and context-dependent effects. Simulations of chromatographic peak integration recapitulated these trends, showing that increased sampling primarily improves integration precision, while quantitative accuracy is strongly governed by peak height and peak shape. Finally, when comparing 20 cancer vs 20 control plasma samples processed with Seer Proteograph, the decrease in DPPP led to a loss of statistical significance for proteins with low-abundance precursors. These findings argue that DIA optimization should prioritize LOQ and statistical power metrics – not identifications alone – by balancing sampling density with chromatographic peak height and quality to maximize useful biological signal.

## Introduction

Modern mass spectrometry (MS)-based proteomics provides quantitative measurements of peptides and their inferred protein groups from complex biological samples, enabling detection of biologically meaningful differences across experimental conditions.^1–3^ Although numerous acquisition strategies have emerged, most rely on broadly similar principles of quantifying chromatographically separated peptide ions and summarizing their associated fragment-ion features. In Data-Independent Acquisition (DIA) MS, peptides are first separated by liquid chromatography (LC) and subsequently isolated in the gas phase using quadrupole mass filtering. Multiple co-eluting precursors of similar mass-to-charge are then co-fragmented and analyzed, yielding multiplexed fragment-ion spectra. Under typical LC conditions, peptide elution profiles exhibit Gaussian-like shapes, and quantitative values are derived through numerical integration of fragment-ion peak areas followed by peptide-level feature summarization.^4^ Similar strategies apply to targeted MS and data-dependent acquisition methods.^5^ Across all methods, achieving high precision and accuracy in peptide quantification is essential; only with sufficiently accurate measurements can biological inference be supported with robust statistical power.^6^

Recent technological advances have expanded the scale and scope of MS proteomics, enabling applications such as single-cell analysis, population-level profiling, and high-throughput translational research.^7–10^ Automated sample handling has improved reproducibility,^11,12^ while higher-flow chromatography and enhanced ion delivery enable robust analysis of hundreds or thousands of injections.^13–15^ Instrumentation improvements–including more sensitive detectors and spectral acquisition rates exceeding 200 Hz–have enabled the use of shorter chromatographic gradients with minimal loss in identifications or reported precision.^16–18^ Strategies to increase identification depth in DIA such as narrow isolation windows have been employed on various instruments.^19–22^ Narrower DIA windows improve specificity and subsequent identifications, but they often reduce sampling density across chromatographic peaks, and the consequences for quantitative accuracy and biological discovery remain under-characterized. Identification depth alone is insufficient for biological inference: studies require accurate, linear quantification that reflects true biological signal rather than peak sampling or integration artifacts. Best-practice guidelines for DIA have suggested a minimum of ten datapoints per peak (DPPP) for accurate quantification.^5,23^ However, the drive toward narrow-window DIA and shorter-gradient, high-throughput workflows has challenged this threshold.^24,25^

Although prior work has proposed DIA method design heuristics, these guidelines rarely quantify how peak sampling strategies affect LOQ yield or the number of statistically powered discoveries in real cohort comparisons. Statistical power is governed by effect size, cohort composition, and observed precision.^26,27^ As depth increases, the required effect size – an observation of quantitative fold change and precision – becomes more stringent. These considerations are exacerbated in population-scale studies, where unbalanced group sizes are common and Cohen’s *d*-based statistical power requirements rise sharply.^28^ When quantification is compromised, even subtle deviations can lead to inflated sample-size demands or, more critically, incorrect biological conclusions.

Here, we systematically evaluate how chromatographic peak sampling influences quantitative accuracy and subsequent differential abundance analysis. Using a matrix-matched calibration curve (MMCC) dataset, we quantify accuracy figures of merit including limit of detection and quantification (LOD, LOQ) of individual precursor ions. We extend this analysis by performing in silico subsampling of DPPP on raw MS files. We complement empirical findings with feature-level modeling and simulation under perturbed chromatographic conditions to characterize determinants of quantification degradation and implications for longitudinal acquisitions. Finally, we apply these insights to an idealized case-control study to demonstrate how sampling of DPPP directly impacts statistical significance and biological discovery. Low DPPP can be effective for increasing proteome coverage, but our results show that reduced sampling may also compromise the fraction of features that remain linearly quantifiable and statistically significant, especially at low abundance. We also observe that DPPP as a performance proxy is insufficient to assess quantitative performance. From these results we develop recommendations for sensitive, high-throughput discovery proteomics experiments to maximize statistical power and improve the accuracy of biological interpretation.

## Methods

### Data Accession

Data for matrix-matched calibration curve (MMCC) experiments were obtained from ProteomeXchange (PXD042704),^16^ hosted via Panorama. Specifically, datasets acquired using 3.5 ms ion injection times and 2 Th precursor isolation widths were used (directory: MMCC_AS_3p5ms_2Th). Chromatogram libraries were constructed using files located in the ASChrLib subdirectory.

Data for the case–control cancer study were retrieved from PRIDE (ProteomeXchange accession PXD060573).^29^ Seer-associated raw files from experiment 5 were selected. Files followed the naming convention `20241014_WFB_exp05_seer_N_NPX.raw`, where N denotes sample indices 1–40 and X denotes nanoparticle type (NPA or NPB). Samples 1–20 corresponded to control samples, and samples 21–40 corresponded to cancer samples.

### File Pre-Processing

All raw data files were converted to mzML format using ProteoWizard MSConvert.^30^ To generate subsampled mzML datasets, custom python scripts were developed to load mzML files into memory and selectively retain MS2 spectra according to precursor isolation window–specific sampling patterns. Sampling schemes were encoded as deterministic binary patterns, where `“1111”` represented full sampling, `“1010”` represented half sampling, and `“1000”` represented quarter sampling. These patterns indicate retention of every spectrum, every other spectrum, or every fourth spectrum, respectively, for a given isolation window. MS1 spectra were not subsampled, as employed MS acquisition does not enforce strict cadence matching between MS1 and MS2 scans. We retained all MS1 scans across conditions; MS2 sampling was isolated to window-specific patterns, so that observed differences reflect MS2 sampling density rather than changes in precursor survey information.

### Search Settings

All data were searched using DIA-NN version 1.8.1,^31^ unless otherwise specified. Newer versions were not used due to licensing constraints. Library prediction of tryptic peptides was done with default GUI specifications in DIA-NN, using: `--gen-spec-lib --predictor.` A combination of human UniProt protein sequences including isoforms and typical MS contaminants was used for prediction (generated 12/9/2022, 71,094 entries). Search command settings were searched as follows: `--qvalue 0.01 --individual-mass-acc --individual-windows –relaxed-prot-inf --smart-profiling --peak-center --no-ifs-removal.` Second-pass search was enabled with the addition of `--reanalyse`. The chromatogram library was generated against the ASChrLib files using predicted library, single-pass search on appropriate mzML files with addition of flag `--out-lib`. DIA-NN was configured to export a library from the search for subsequent chromatogram library–based analyses with addition of flag `--lib`. Searches using chromatogram libraries were performed in single-pass mode. For predicted library workflows, the initial search is referred to as “Predicted”, while “Predicted, MBR” or “MBR” refers to the second pass, match between runs, search following construction of an empirical library incorporating observed retention times and relative fragment ion intensities.

### Consensus Peak Area Estimate

For each search strategy (chromatogram library, predicted library, and MBR), precursor intensities were summarized independently. Within each search strategy, precursor peak area was evaluated at the 100% concentration condition. Replicate measurements were summarized as the median of `Precursor.Normalised` values to reduce sensitivity to outliers and replicate-level variability. This per-strategy peak area summary was used for all downstream analyses, including stratification by peak area quartile and modeling of figures of merit. No averaging or reconciliation of intensities across search strategies was performed. Within each search method, precursors were binned into four equally sized quartiles based on consensus peak area estimations. Quartile assignments were used to stratify analyses and control for dynamic range effects.

### Datapoint Per Peak Estimate

Because DIA-NN does not directly report datapoints per chromatographic peak (DPPP) in output, DPPP was estimated using a peak-width–based heuristic derived from DIA-NN version 1.8.0 operated in “High-Accuracy, Any LC” quantification mode. For each precursor observation, peak duration was calculated from the reported retention time boundaries (`RT.Stop` *−* `RT.Start`). We inferred an effective cycle time from peak-width-derived sampling and calibrated it using the narrowest observed peaks as a conservative lower-bound (4 points per peak in DIA-NN report), providing a calibrated estimate for sampling intervals.

Precursors were separated into two classes based on their reported retention time boundaries relative to cycle time. For precursors exhibiting ≥5 estimated datapoints per peak, DPPP values were evaluated at the highest concentration level available (typically the undiluted condition), summarized across replicates using the median, and rounded to the nearest integer. For precursors with fewer than five datapoints per peak, DPPP could not be reliably inferred from reported peak width alone. In these cases, a resampling-based heuristic was applied using multiple MS2 retention patterns representing all possible single-window and multi-window sampling configurations corresponding to 1–4 datapoints per peak. The pattern used for two-DPPP included `1100`, `0110`, `0011`, and `1001` such that only patterns with consecutive scan retention were evaluated. For each precursor, the frequency of identification across these sampling patterns was weighted by the number of datapoints retained, and the mean weighted score across replicates was rounded up to the nearest integer and capped at four datapoints per peak.

For downstream stratification we assigned each precursor a consensus DPPP equal to the maximum estimate observed across workflows, reflecting the best-supported sampling density observed for that feature across analyses. The resulting precursor-level DPPP distribution was approximately normal, with >99.7% of precursors identified at 1% FDR in DIA-NN 1.8.1 also identified at 5% FDR in DIA-NN 1.8.0. Precursors without DPPP annotated were excluded from DPPP-dependent analyses.

### Figures of Merit Calculation

Limits of detection (LOD) and quantification (LOQ) were calculated using an implementation available on GitHub (https://github.com/lindsaypino/matrix-matched_calcurves/tree/master/bin, `calculate-loq.py` commit 959f6d1). DIA-NN `report.tsv` outputs were converted to EncyclopeDIA-like, precursor-centric input formats prior to analysis. The script was executed with the `--min-noise-points 0` parameter to accommodate model assumptions and performance differences between DIA-NN and EncyclopeDIA. DIA-NN reports filter on local *q*-value unlike EncyclopeDIA, thereby not returning as many low but non-zero peak area values that are otherwise used as input into LOD setting. Resulting figures of merit are provided in the uploaded results.

### Linear Modeling of Datapoints Per Peak and Peak area versus Figures of Merit

To quantify how DPPP and precursor peak area influence analytical sensitivity, we modeled the response of log_10_ LOD or LOQ using linear regression. Predictor variables precursor peak and DPPP were modeled on a log_2_ scale so that coefficients represent the effect of doubling. To capture nonuniform behavior across the dynamic range of intensities, precursors were stratified into peak area quartiles (peak area_q_), and interactions were included to allow slopes to vary by peak area quartile. The full R glm formula was specified as

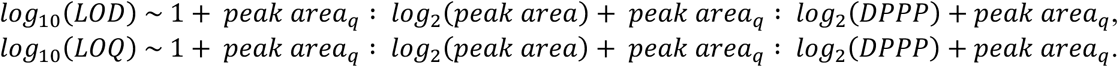

Model coefficients (*β*) are estimated on the log_10_ scale and represent additive changes in log_10_ LOD or LOQ per doubling of predictor. For interpretability, coefficients were transformed to percent change in LOD or LOQ using (10^%^ − 1) × 100%. Confidence intervals were computed as Wald intervals on the log_10_ scale and transformed identically.

### Association of Feature Analysis

To visualize non-linear associations between analytical performance metrics, generalized additive models (GAMs) were fit using `mgcv` (v1.9-4) in R. For each relationship, the model was specified as

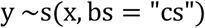

where *y* is the response variable, *x* is the predictor, and *s(·)* denotes a smooth term modeled using a cubic regression spline basis with shrinkage penalty *(bs = “cs”)*. Smoothing parameters were estimated by restricted maximum likelihood. To improve robustness to heavy tails and heteroscedasticity, models were fit using the scaled t-distribution (family = “scat”). Trend lines are displayed as fitted smooths *ŷ(x)*. Effective degrees of freedom (edf) and deviance explained (dev.expl) were extracted as descriptive measures of relationship complexity and association strength.

### Simulation of Quantification

To evaluate the effect of sampling density on chromatographic peak integration accuracy, simulations of extracted ion chromatograms were performed using an exponentially modified Gaussian peak model.^32^ Peak shapes were parameterized by a Gaussian width and an exponential tailing parameter derived from a discrete symmetry factor to model increasing chromatographic tailing. All simulated peaks were normalized such that the true integrated area equaled 1, enabling direct quantification of integration error. Peak area measurements were sampled uniformly across each peak using a fixed number of DPPP with widths determined by elution profile.

Multiple sources of experimental variability were incorporated to reflect realistic LC-MS acquisition conditions. A time-correlated baseline was simulated using an autoregressive process, with mean baseline proportional to the peak height, a defined 5% coefficient of variation controlling baseline fluctuation magnitude, and a correlation parameter governing baseline smoothness. This establishes the bounds of peak integration.

Random normal noise was additionally added with 5% standard deviation of absolute observation modulation specified. Negative intensities were truncated to zero. Peak areas were estimated using a trapezoidal integration method and error was assessed as absolute percent deviation from ground-truth area, i.e., 1. Assigned inclusion of variance are arbitrary metrics not necessarily corresponding to existing MS instruments.

Simulations were performed across a grid of DPPP, SNR and peak symmetry conditions with 500 replicate peaks per parameter condition. Additional longitudinal simulations were conducted to model performance drift over time under scenarios of progressive chromatographic broadening, declining SNR, or both, while holding baseline characteristics constant. All simulations were independent.

### Cancer Study Statistical Analysis

Files from the cancer cohort were searched in DIA-NN (v1.8.1) with the predicted library, MBR approach. Raw files were analyzed for each of two nanoparticle (NP) wells (n=40 samples, n=80 files), with samples 1–20 corresponding to healthy controls and 21–40 corresponding to cancer. Cohort data were searched using the MBR method, filtered at 1% *q*-value on local, library, and library protein group levels, and NP-precursors were filtered at 50% completeness within groups and required to be present in both groups prior to differential testing. Precursors were annotated as nanoparticle-precursor (NP-precursor) identifications per fraction as well as sample group based on run metadata. Within sample groups, NP-precursors were filtered at 50% completeness and again by presence in both cancer and healthy conditions.

Two separate search analyses were done: quantile-sensitive analysis of unmodified mzML files and peak sampling completeness analysis at quarter, half and full sampling levels. For the quartile-stratified analysis, quartiles were defined after completeness filtering to equalize comparison set size. For each precursor, a feature-level peak summary was computed as the mean reported peak area across all samples and were then ranked by this summary into four equal bins.

For each NP-precursor in each respective analysis, differential abundance between Cancer and Healthy groups was assessed with limma’s empirical Bayes moderation on DIA-NN reported normalized intensities yielding moderated t-statistics and two-sided *p*-values for each precursor. Multiple hypothesis testing was controlled using the Benjamini-Hochberg (BH) false discovery rate procedure. For the peak sampling completeness comparison, each peak sampled dataset was treated as an independent hypothesis family and BH correction was applied within each dataset. For the quartile-stratified analysis, BH correction was applied globally across all precursors in that dataset, with results subsequently summarized and visualized by quartile. Statistical significance was defined with BH-adjusted *q-*value threshold 0.05.

To quantify the detectability of observed group differences, achieved statistical power was computed per precursor using a standardized effect size derived from the limma model. Specifically, observed effect size was defined as

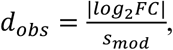

where |log_2_FC| is the absolute limma estimated log_2_ fold-change and *s_mod_* is the limma moderated residual standard deviation. Sample sizes were defined per precursor with *n_H_* and *n_C_* defined as number of non-missing observations in Healthy and Cancer groups, respectively. Achieved power was computed using a two-sample t-test power function with unequal sample sizes, observed effect size and two-sided significance level α = 0.05. Precursors were classified as “Powered” if they achieved at least 0.80 power.

Because BH correction was performed within each peak sampling completeness condition, significance counts reflect within-condition discovery control rather than a shared hypothesis family; comparisons therefore reflect a combined effect of detection and quantification changes under each condition.

## Results

To evaluate determinants of quantitative performance in DIA proteomics, we analyzed a publicly available matrix-matched calibration curve (MMCC) dataset acquired on a ThermoFisher Orbitrap™ Astral™ MS.^16^ Quantitative performance was evaluated for key figures of merit: limit of detection (LOD) and limit of quantification.^23^ LOD reflects detectability above a background noise, whereas LOQ represents the lowest concentration at which a precursor maintains an approximately linear response while meeting a CV ≤ 20% criterion under bootstrap resampling of observations and their respective linear models.

To isolate the effect of chromatographic sampling on quantification outcomes, we modulated peak sampling completeness by deterministically thinning out MS2 spectra. It was done at full, half, and quarter patterns, where the respective fraction of equally spaced MS2 spectra per isolation window were retained in retention-time resolved pattern (e.g., “half” refers to every other MS2 spectra being retained for full acquisition time). All peak sampling completeness conditions were searched under three search approaches: chromatogram library, predicted library, and predicted library with second-pass MBR search (“predicted, MBR” or “MBR”).

### Qualitative Metrics

In MMCC experiments, unlabeled samples are diluted by an isotopically labeled pair or similar matrix sample. Consistent with this design, precursor identifications decreased with decreasing concentration of target matrix (Fig. 1B; Supp. Fig. S1). In the chromatogram library search, 130,498 mean precursors were observed at 100% concentration whereas 52,768 were measured at 10% concentration. Across all library workflows, reduced peak sampling completeness decreased the number of observed precursors, consistent with reduced chromatographic evidence supporting 1% *q*-value identification thresholds. The half and quarter peak sampling datasets decreased to 118,051 and 84,901 observed precursors at 100% concentration, respectively.

**Figure 1.**
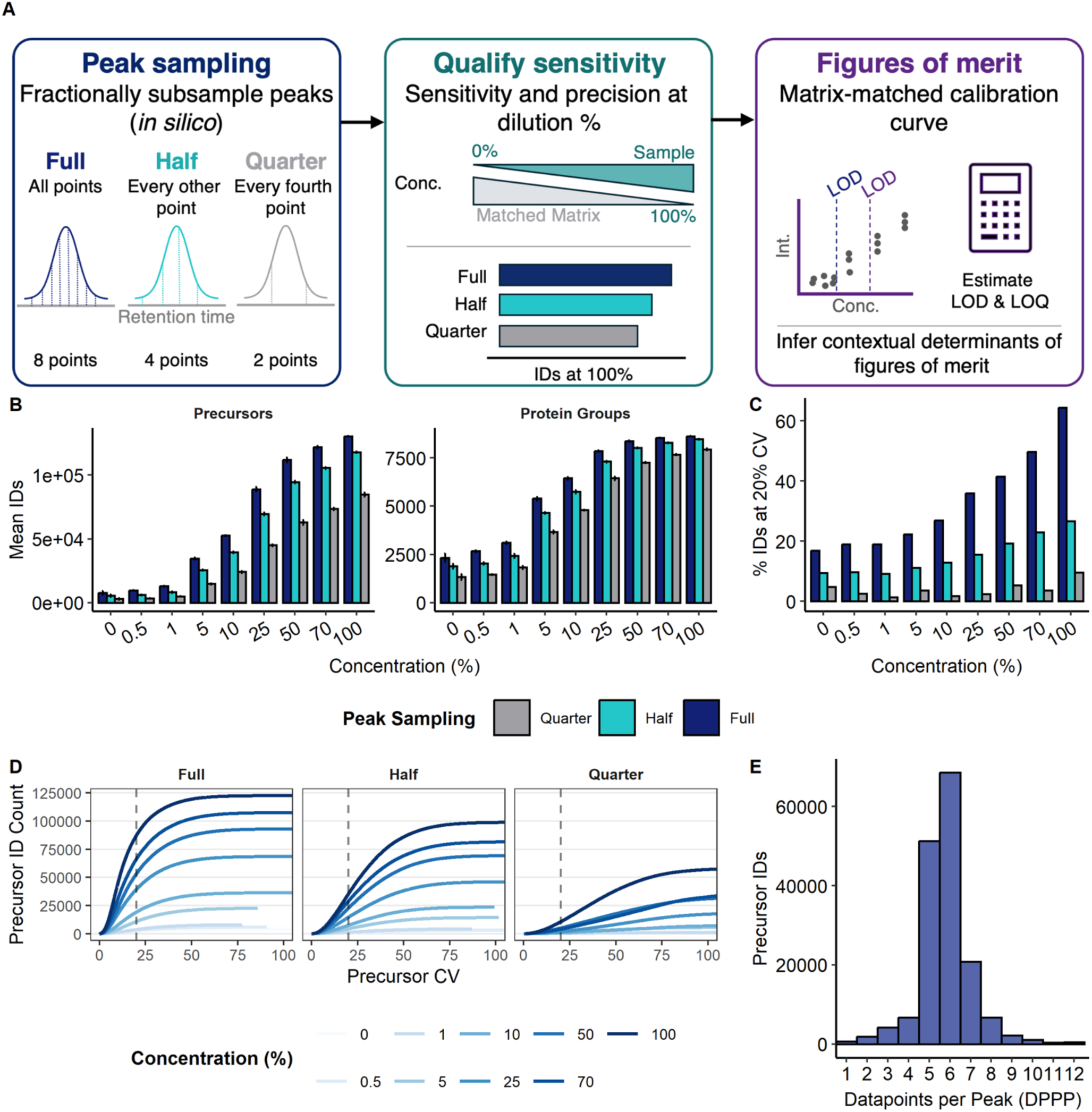
– (A) Study overview. Peak sampling methodology before sensitivity and figure of merit calculation. (B) Mean precursor and protein-group identifications across matrix-matched calibration curve concentrations under full, half, and quarter MS2 sampling. (C) Fraction of identified precursors with CV ≤ 20% at each concentration for full/half/quarter sampling. (D) Precision accumulation curves showing the number of precursors achieving CV specified on the x-axis. (E) Distribution of DPPP estimates for precursors retained for downstream analysis. Figures from Chromatography Library search.

Reduced peak sampling completeness also degraded precision distributions. Across library strategies, precision curves were systematically depressed as peak sampling completeness declined (Fig. 1D; Supp. Fig. S2), and fewer precursors fractionally met a 20% CV threshold under half and quarter completeness (Fig. 1C; Supp. Fig. S3). At 100% concentration in chromatogram library search, 64.3, 26.7, and 9.6% of precursors achieved this threshold for full, half, and quarter peak sampling respectively. Chromatogram library searches retained the strongest fractional precision performance across peak sampling completeness conditions, while predicted and MBR workflows observed weaker accumulative performance by comparison (Supp. Fig. S2). To contextualize additional differences, precursor peak areas were computed per workflow using the highest concentration where a precursor was observed and summarizing replicates as the median search engine normalized values. The total count of observed precursors, not median, was approximately 14% greater for either predicted method than chromatogram method. However, 90% of precursors were measured in all three replicates of the chromatogram library method, while 70% and 88% were measured as complete in the predicted and MBR datasets, indicating benefit of experiment-specific libraries for reproducible identification. 12% more precursors were measured with completeness in 3 replicates at 100% concentration in MBR than in the chromatogram library indicating benefit of broad scope initial search before library refinement. Heuristic estimate of chromatographic sampling per precursor yielded a DPPP distribution spanning 1–12 with a mode near 6 in the full dataset at undiluted concentration, not achieving benchmark recommendations of 10 mean DPPP (Fig. 1E).

### Empirical Figure of Merit Observation

We next evaluated LOD and LOQ in the MMCC curve using a protocol previously described.^23^ In this approach, each figure of merit is reported as the fractional concentration of the endogenous precursor signal at which the LOD or LOQ criterion is satisfied. Precursors that do not meet the criterion at any tested concentration are assigned an infinite estimate. Across peak sampling completeness conditions, LOD distributions closely tracked identification losses (Fig. 2A–B; Supp. Fig. S4–S5), indicating that reduced sampling primarily removes features rather than shifting detectability for those that remain detected and that DIA-NN has more limited reporting of sub-noise background peak area. In contrast, LOQ degraded markedly with reduced peak sampling completeness: nearly 80% of precursors in the full condition had observable LOQ, whereas fewer than 25% remained quantifiable under quarter completeness, with half completeness yielding an intermediate effect (Fig. 2A–B). These patterns persisted under cohort-relevant LOQ stringency (fractional LOQ ≤0.50, corresponding to ≥2-fold quantifiable change), showing that reduced sampling increases both the proportion of precursors with infinite LOQ and the fraction whose LOQ exceeds biologically useful concentrations. Trends were also consistent across library paradigms, indicating that peak sampling completeness impacts quantifiability independent of search strategy.

**Figure 2.**
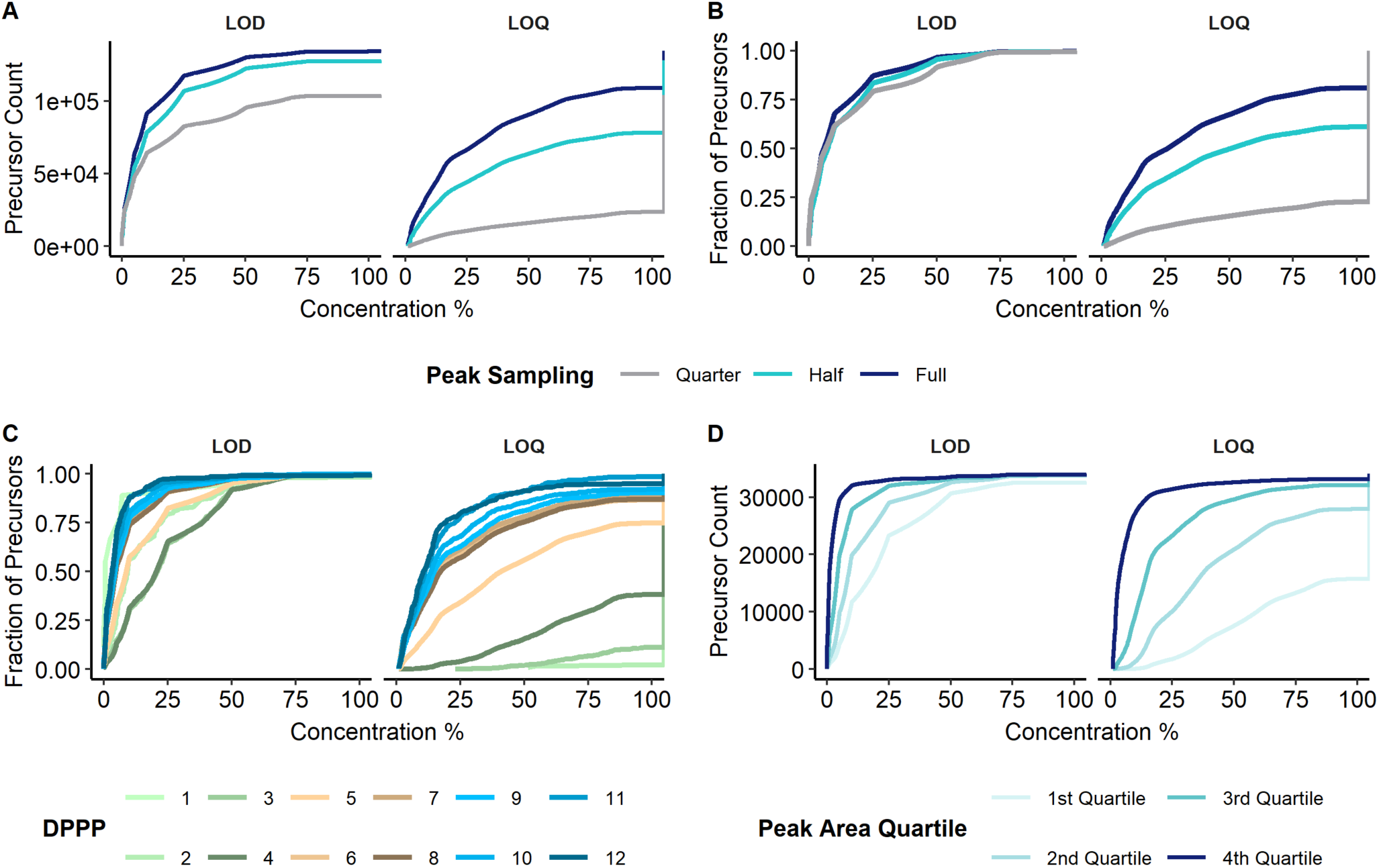
– (A) Accumulation curves showing the number of precursors with LOD (left) or LOQ (right) at or below a given fractional concentration threshold in the calibration series, for full/half/quarter MS2 sampling (chromatogram-library). (B) Fraction of precursors at LOD and LOQ values for the same data in (A). Precursors that never meet the LOD or LOQ criterion across tested concentrations are assigned infinite values and retained. (C) Fraction of precursors at specified LOD or LOQ according to DPPP estimation. (D) Precursor figure of merit distributions stratified by precursor area quartile.

To evaluate whether DPPP alone explains LOQ behavior, precursors were evaluated by DPPP estimates at 100% concentration (Fig. 2C; Supp. Fig. S6–S9). Precursors with <5 DPPP exhibited markedly poorer LOQ performance, whereas precursors achieving ≥6 DPPP exhibited progressively improved LOQ behavior with diminishing returns at higher DPPP. Although LOQ improved with increasing DPPP, the separation was incomplete, motivating evaluation of additional contextual determinants such as peak area and peak-shape effects.

Data stratification by peak area quartile at 100% concentration further revealed stronger differentiation in LOQ performance by DPPP group alone (Fig. 2D; Supp. Fig. S10). Generalized additive modeling (GAM) of feature associations supported these observations, with LOQ showing the strongest association with peak area and weaker deviance explained for DPPP comparisons (Supp. Fig. S11–S18; Table 1). Because DPPP is discretized and estimated, deviance explained for DPPP associations should be interpreted more descriptively than as a definitive measure of association strength. Weaker deviance explained is therefore anticipated for DPPP-dependent analyses.

**Table 1.**
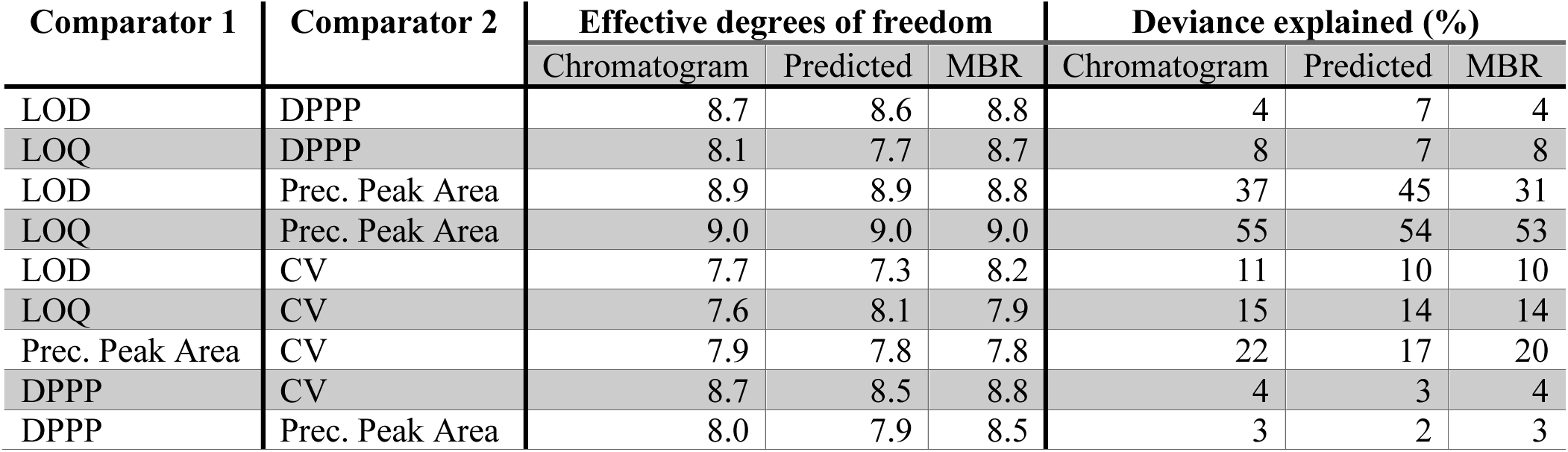
Nonlinear feature associations with quantitative performance metrics across workflows. GAMs were fit for each comparator pair within each search strategy. E[ective degrees of freedom (edf) summarize relationship flexibility (higher values indicate more complex nonlinearity), and deviance explained quantifies the strength of association between variables (higher values indicate better fit). “Prec. Peak Area” denotes precursor peak area at 100% concentration.

LOQ losses under peak sampling were consistent across all spectral library methods, indicating that reduced peak sampling affects quantifiability relative to detectability regardless of search paradigm. However, library differences impacted the distribution of quantifiable features with respect to estimated DPPP. Workflows that expanded coverage to lower-abundance precursors (predicted and MBR) tended to exhibit broader LOQ distributions in each DPPP boxplot. Chromatogram libraries also observed more monotonic relationships of DPPP to median CV and precursor peak area, whereas more complex relationships were observed for other methods.

Taken together, these observations support a context-dependent model of quantifiability best-practice as opposed to singular metric proxies.

### Joint Modeling Supports Context-Dependent EFects of Sampling Density

To quantify joint contributions of peak area and DPPP, we modeled log-transformed LOD and LOQ as functions of log_2_ peak area and log_2_ DPPP, stratified by peak area quartile at 100% concentration. Across quartiles and library paradigms, increased precursor abundance consistently improved both LOD and LOQ. In contrast, DPPP effects were inconsistent and often neutral or unfavorable (Fig. 3). A plausible interpretation is that peaks with higher DPPP may reflect broader or more tailing peak shape that distribute signal over longer timescales, thereby reducing peak height above baseline and complicating integration, while additional sampling is most beneficial in low-abundance regimes where peak focusing is not observed. The magnitude of signed effect is often greater for peak area than DPPP, however the Wald uncertainty intervals show greater interval for DPPP than peak area, perhaps as a limitation of more discretized observation range. Peak height may then be a stronger predictor of quantification performance than simple peak area. Together, linear modeling and GAM analyses support assertion of peak area as a stronger associator to quantification accuracy, whereas DPPP has weaker and context-dependent effects.

**Figure 3.**
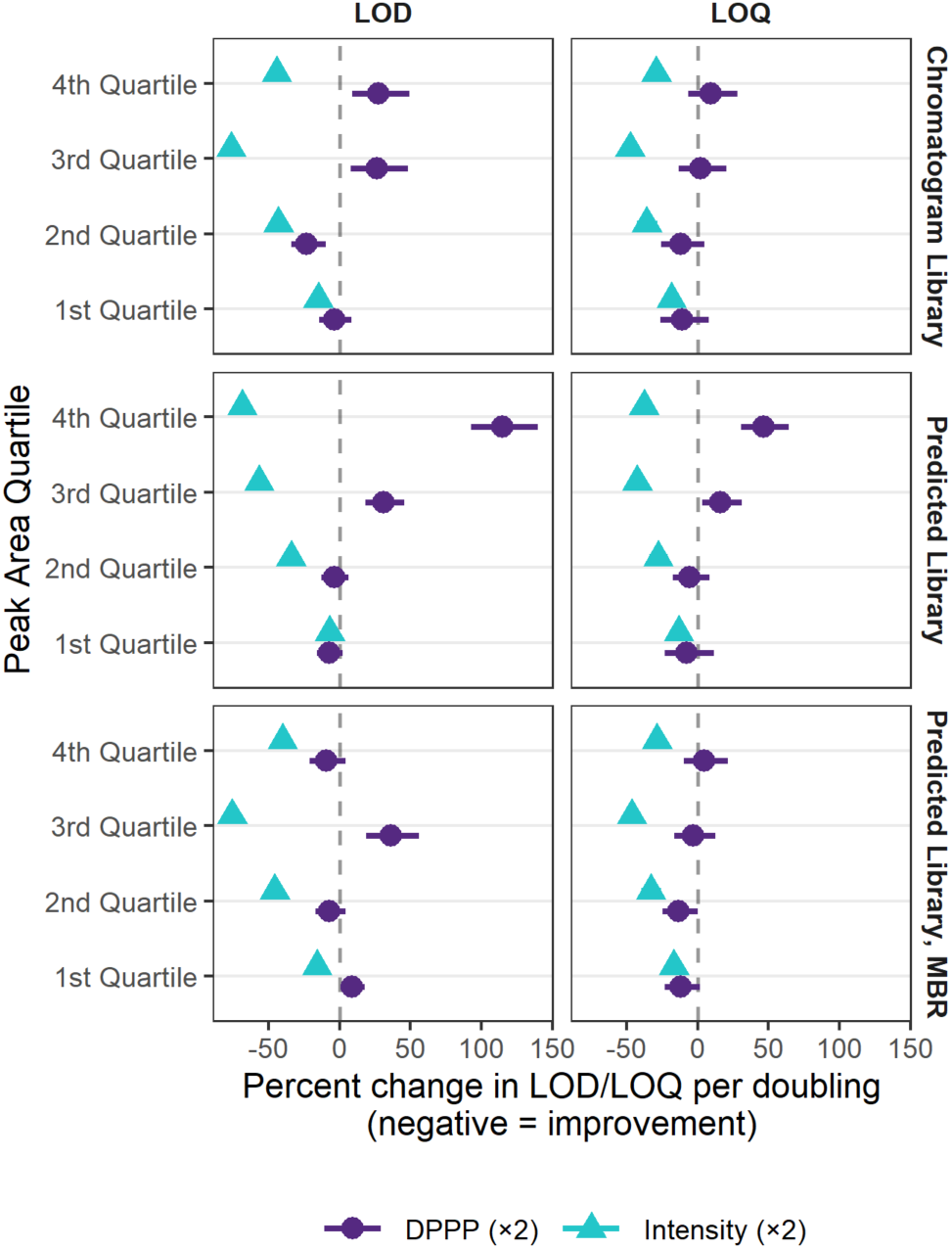
– (A) Linear model coefficients estimating percent change in log_10_(LOD) or log_10_(LOQ) associated with either doubling precursor peak area (log_2_ scale) or doubling sampling density (DPPP; log_2_ scale), stratified by peak area quartile and search. Negative values indicate improved sensitivity (lower LOD/LOQ). Points indicate model estimates and error bars indicate Wald confidence intervals.

### Simulated Accuracy and Precision

To further contextualize these trends, simulations were performed using exponentially modified Gaussian peaks above a constant noise background across a grid of DPPP, signal-to-noise ratio (SNR; proxy for interpretable peak height above noise background), and peak symmetry (Fig. 4). Peak area was estimated by trapezoidal integration and compared to a unit ground-truth area using absolute percent error. Across simulated conditions, median bias approached zero between 3 and 4 DPPP. However, increasing DPPP beyond 4 did not uniformly reduce bias, plateauing with values on interval (-0.03, 0.03) across much of the SNR and symmetry grid, consistent with persistent baseline and stochastic noise contributions. Peak symmetry and SNR did not significantly modulate bias.

**Figure 4.**
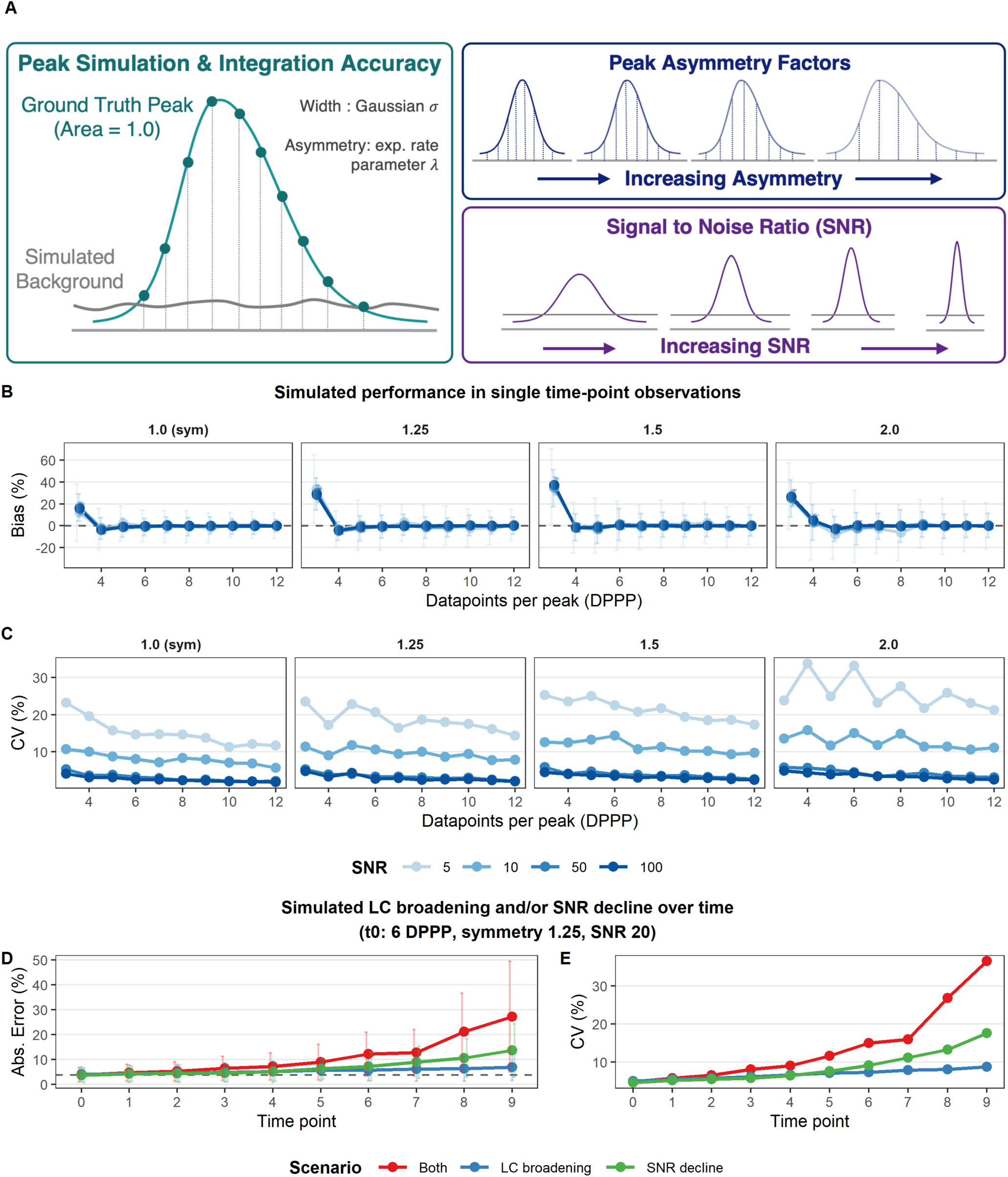
(A) Summarization of simulation characteristics for peak determination and impact of parameterization on peakshape. Bias of simulated peaks under various sampling frequencies and asymmetry parameterization. Error bars are standard deviation of measure. (C) Integration precision for the same conditions as in (A). (D) Longitudinal simulation of increasing absolute percent error over time under progressive chromatographic broadening, declining SNR, or both (baseline condition: DPPP=6, symmetry=1.25, SNR=20). (E) Corresponding longitudinal increase in CV.

Integration precision, measured by CV across replicate realizations, improved with increasing DPPP and SNR. Trends were consistent across different SNR and peak asymmetries, however, greater proportional improvements to precision were observed with increasing DPPP at high SNR. Greatest decrease in CV was observed for low SNR, high peak asymmetry conditions. The quantification robustness improvement may then reflect dependence on scan placement within a chromatographic elution. Because precision contributes directly to LOQ evaluation, DPPP-influenced CV improvements are expected to most strongly reduce LOQ under low-abundance or high peak asymmetry conditions.

To simulate longitudinal chromatographic degradation, we modeled progressive twofold peak broadening coupled to a fourfold SNR loss (Fig. 4D-E). Under these degradation conditions, median absolute error increased from 4.2% to 27.4%, and CV increased from 3.2% to 20.1%, sufficient in some regimes to shift measurements beyond LOQ criteria (typically <20% CV). Collectively, these simulations support a model in which DPPP primarily modulates integration variability, while peak context – particularly peak height (SNR) and peak shape – most strongly governs accuracy and secondarily contributes to precision. Consistent with this interpretation, we distinguish integrated peak area from peak height as related but non-equivalent measures. Although our empirical analyses focus on integrated peak area, peak area is only an indirect proxy for the effective signal above the instrument noise background, mediated by the co-occurring peak shape and peak width. While these results indicate that detectable peak height above the noise background (hereafter “peak height”) is mechanistically central to quantitative performance, we do not directly quantify peak height in this study. Standard DIA-NN outputs report integrated quantities, and peak-height-based quantification remains an experimental mode; accordingly, peak height is discussed here as an inferred driver supported by simulations of peak-context modeling.

Precision in this simulation reflects detector noise and stochastic scan placement; additional variance from chromatography and pre-analytical factors would further inflate observed CV in real studies. A simple case–control power simulation (Supp. Fig. S19) demonstrated that increasing CV reduces detection power for a fixed effect size, requiring either reduced statistical power thresholds or otherwise requiring significant increases in cohort sizes. These recommended cohort size increases may be practical with higher throughput observation but requires availability of sufficient samples.

### Application to Biological Cohorts

To evaluate how peak sampling propagates to biological inference, we applied the peak subsampling strategy to a publicly available 20 vs. 20 cancer versus healthy blood plasma cohort generated using the Seer Proteograph™ XT™ workflow on an Orbitrap Astral MS. This analysis enabled assessment of how reduced datapoints per peak affects differential-abundance testing, multiple-testing corrected significance, and achieved statistical power (Fig. 5).

**Figure 5.**
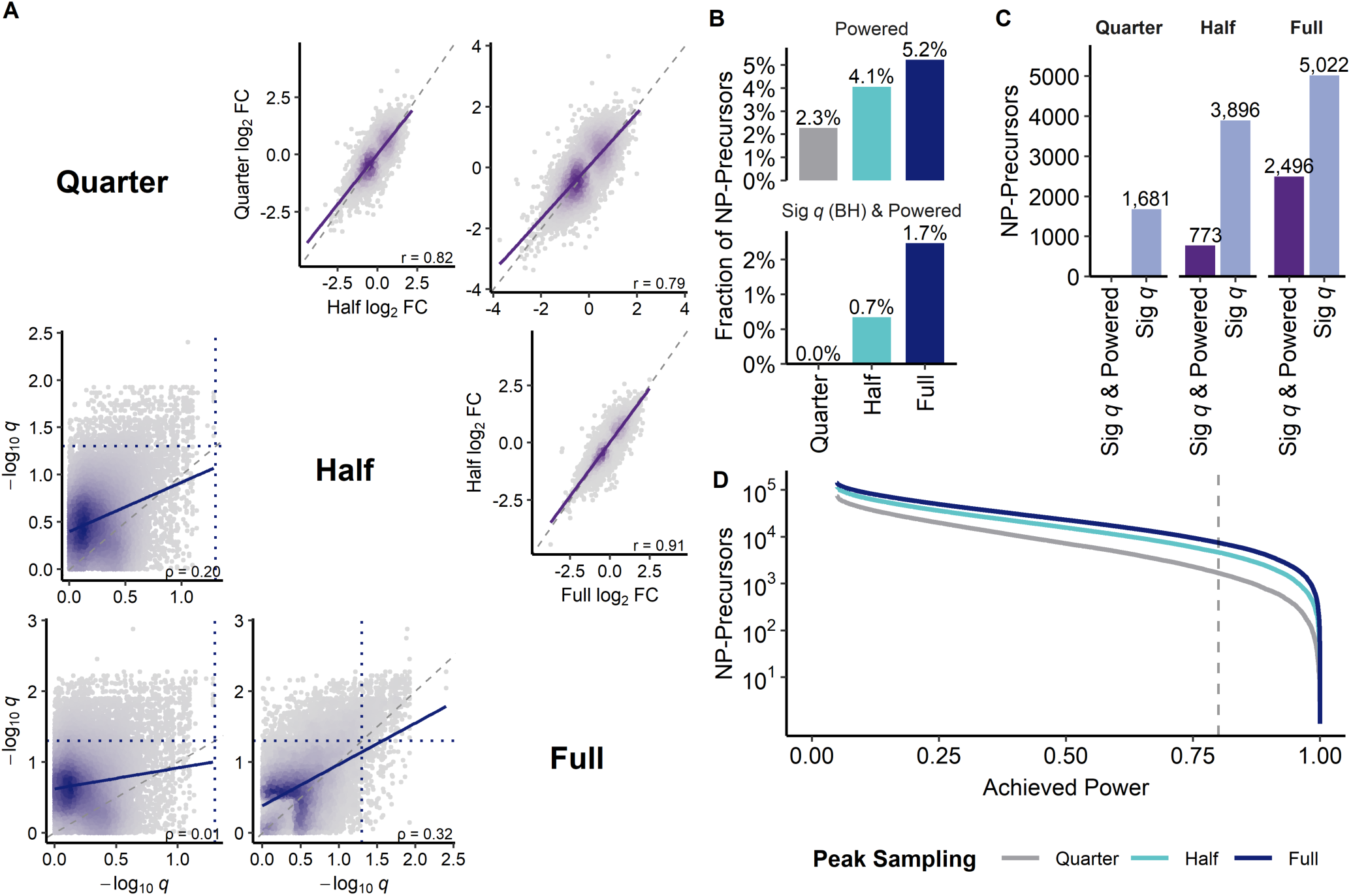
– Evaluation of statistical signal in a 20 vs 20 cancer vs healthy cohort. Significance set at Benjamini–Hochberg adjusted q ≤ 0.05 power threshold ≥0.80 when relevant. (A) Pairs plots of precursors observed in at least one dataset with p-values ≤ 0.05. Top triangle set demonstrates pairwise comparison of shared precursors FC, Pearson correlation and unweighted linear regression. Bottom triangle demonstrates q-value distribution with 0.05 significance threshold indicated. Additional Spearman rho and unweighted linear regression are shown. Identity slope lines included in each half triangle set. (B) Fraction of NP-precursors classified as powered and as both powered and BH-significant across sampling conditions. (C) Total counts of BH-significant NP-precursors and BH-significant + powered NP-precursors for each sampling condition. (D) Accumulation curves of achieved power across NP-precursors, with the 0.80 threshold indicated.

Across peak sampling conditions, differential-abundance structure was broadly preserved, but reduced peak sampling substantially decreased the number of statistically significant and adequately powered discoveries (Fig. 5B–D; Supp. Fig. 20). Pairwise comparison of NP-precursors with nominal evidence of differential abundance in at least one sampling condition, defined as *p* < 0.05, revealed distinct behavior for *q*-value and fold-change concordance (Fig. 5A). In the lower triangle, −log10(*q*) values were only modestly correlated across sampling conditions, with weaker agreement as sampling conditions diverged. This indicates that reduced peak sampling alters statistical significance after multiple-testing correction, likely through combined effects on missingness, quantification precision, and feature-level variance. In contrast, the upper triangle showed stronger concordance in cancer versus healthy log2 fold-changes, particularly between adjacent sampling conditions.

However, comparisons involving Quarter sampling showed greater deviation from the identity line, indicating that reduced datapoints per peak can still alter quantitative effect estimates even when the overall direction and rank structure are partially preserved.

The fraction of powered NP-precursors increased from 2.3% in Quarter sampling to 4.1% in Half and 5.2% in Full (Fig. 5B). More stringently, the fraction of NP-precursors that were both significant after BH correction, *q* ≤ 0.05, and adequately powered, power ≥ 0.80, increased from 0.0% in Quarter to 0.7% in Half and 1.7% in Full (Fig. 5B). In absolute terms, significant NP-precursors increased from 1,681 in Quarter to 3,896 in Half and 5,022 in Full, while significant-and-powered precursors increased from 0 in Quarter to 773 in Half and 2,496 in Full (Fig. 5C). These trends were consistent across the full achieved-power distributions, with Full sampling retaining more NP-precursors at high power thresholds (Fig. 5D). Because BH correction was performed independently within each peak sampling condition, these significance counts reflect condition-specific discovery control rather than a shared hypothesis family; comparisons therefore capture the combined effects of altered missingness, quantification precision, and multiple-testing behavior under each sampling condition.

To contextualize these effects across precursor abundance, we stratified the Full sampling dataset into peak area quartiles. Precursors meeting at least 50% completeness in both cancer and healthy samples were retained, enabling comparison across equal-sized abundance strata (Supp. Fig. S21). The number of significant and statistically powered NP-precursors increased monotonically across peak area quartiles, from 229 in the lowest abundance quartile to 1,811 in the highest (Supp. Fig. S21C), indicating that retained biological signal was enriched among higher-abundance precursors. Together, these results show that reduced peak sampling diminishes recovery of significant, well-powered biological differences in a real case–control setting, and that lower-abundance precursors are less likely to be detected as both differentially abundant and adequately powered.

Although a causal relationship cannot be established without ground-truth abundance, the observed dependence of power on sampling, precision, and abundance is consistent with the reduced precision measured for subsampled peaks and low-abundance precursors in the MMCC dataset (Fig. 1B; Supp. Fig. S19).

## Discussion

Momentum in MS-based proteomics has increasingly favored high-throughput acquisition strategies that preserve robust operation while expanding quantitative depth, enabling studies at cohort scale. These gains have been driven by instrumentation advances that support faster scanning and improved ion detection, making shorter gradients and more aggressive DIA schemes increasingly practical. In parallel, analytical method development is frequently guided by proxy metrics – such as precursor identifications, peptide counts, precision, and DPPP – that are useful for method tracking but can be insufficient to predict whether quantifiable content and statistical power are preserved under design changes. Together, these results argue that no single proxy metric is sufficient: sampling density, peak height, and peak shape jointly determine whether increased nominal depth translates into quantitatively reliable – and biologically powered – measurements.

Across the MMCC experiment, peak sampling minimally altered LOD distributions yet substantially degraded LOQ, indicating that detection can remain largely intact while quantitative fidelity declines. Empirical modeling and simulation helped explain this separation. Precursor signal context – particularly effective peak height relative to local noise/interference and peak shape – dominated integration accuracy and sustained precision. By contrast, DPPP primarily influenced integration precision, and its impact was more context dependent. Case versus control study application translated these quantitative effects into biological outcomes: reduced sampling completeness preserved overall differential-abundance structure but markedly decreased the number of statistically significant and well-powered discoveries. Under reduced sampling completeness, the significant and well-powered discoveries that remained were enriched for higher-abundance precursors, consistent with quantification becoming increasingly constrained by peak context rather than sampling density alone. Together, these findings reinforce that identification depth alone is insufficient to predict cohort-scale biological yield and that method evaluation should emphasize quantifiable content and statistical utility.

Collectively, these findings explain why acquisition changes that increase nominal depth can still reduce biological utility: quantifiable signal depends on both sampling density and peak context, and LOQ losses translate directly into fewer powered discoveries. Gradient shortening compresses chromatographic peaks relative to the acquisition cycle, effectively reducing sampling completeness and increasing sensitivity to scan placement within the peak. Peak compression may also improve peak height, and without decline in peak symmetry may preserve quantifiability even under constrained sampling (*e.g*., lower DPPP). Likewise, increasing the number of DIA windows per cycle can reduce sampling density unless scan speed is correspondingly improved. Our results suggest these strategies may increase nominal depth while decreasing the fraction of features that remain reliably quantifiable and statistically actionable, particularly for low-abundance precursors. Because statistical power in cohort studies depends jointly on effect size, measurement variance, and feature count, throughput-oriented methods without coupled increase in cohort size may enable in-depth identifications at the expense of quantification fidelity and may paradoxically reduce powered discoveries.

Beyond sampling and window design, instrument method choices such as restricting the MS2 scan range to concentrate acquisition on highly populated m/z regions may increase protein group counts in some settings but can reduce sequence coverage and limit detection of informative peptides. In population-scale designs, this may carry downstream costs for analyses that depend on peptide-level diversity, including pQTL discovery and proteoform-aware inference. Method optimization should also be tailored to biosample composition and analytical goals. Blood plasma, for instance, spans more than 8 orders of magnitude in dynamic range,^33^ with a small number of highly abundant proteins and a long tail of low-abundance species. In this context, prioritizing datapoints per chromatographic peak may improve quantitative robustness and power for differential abundance analyses, even if total identifications are constrained. By contrast, alternative matrices may exhibit a compressed dynamic range or greater representation of mid- to high-abundance proteins, supporting higher-throughput acquisition while maintaining strong quantitative yield. Across matrices, optimal method selection should also account for measurement objectives, such as reliable quantification of low-intensity biomarkers, where modest reductions in throughput may be justified to sustain quantification of high value targets

For cohort-scale discovery proteomics where the objective is maximizing statistical yield, we highlight the following practical considerations building on these results and existing best practices. (i) Use sample-aware acquisition strategies that maximize statistically actionable features rather than nominal depth, recognizing that biosample matrices differ substantially in abundance dynamic range (*e.g.,* cell lysate versus plasma) and therefore impose different constraints on the peak context regime required for reliable quantification. (ii) Benchmark methods using LOQ yield and precision (e.g., fraction of features with finite LOQ and CV ≤20%) alongside identification depth, since nominal depth alone does not determine the set of statistically actionable measurements. (iii) In place of per-precursor monitoring that is difficult to standardize across biosamples, use standardized peptide mixtures spiked into study samples to monitor peak shape and effective points-per-peak across abundance regimes. Continuous monitoring through toolkits like Skyline and Panorama^34^ serve as practical ways to execute quality assurance. (iv) When LOQ degrades following a method modification, our results suggest first diagnosing whether reduced peak height or reduced sampling density is the primary driver, and prioritizing recovery of the limiting factor before further increasing scan burden, while also considering that detector saturation can degrade LOQ at the upper end of the dynamic range. (v) Track standard peptide peak width and asymmetry as quality control metrics throughout study duration and intervene when systematic peak broadening or tailing is observed. While non-exhaustive, these recommendations support a contextual framework in which sampling density, peak height, and peak shape must be jointly considered to preserve quantification fidelity and statistical power in high-throughput DIA proteomics, enabling more accurate and reproducible biological inference in cohort-scale studies.

## Conclusion

In this study, we systematically evaluated how chromatographic sampling influences quantitative performance in DIA proteomics using MMCC-derived figures of merit and controlled modulation of peak completeness. Reduced peak completeness minimally affected detectability (LOD) but substantially degraded quantifiability (LOQ), indicating that identification-centric readouts can remain stable while quantitative utility declines. Empirical modeling and peak-integration simulations further showed that peak area and shape are stronger determinants of sustained quantifiability than datapoints per peak alone, with reduced sampling primarily worsening integration variability. Finally, application to a 20 vs 20 case–control cohort demonstrated that decreased peak sampling completeness reduces the number of statistically significant and powered discoveries, with the retained discoveries enriched for higher-abundance precursors. Together, these results emphasize that acquisition strategies for large-scale discovery proteomics should be evaluated by their ability to preserve contextual quantification quality – balancing throughput, MS method design, and sample loading to maintain sufficient sampling and peak height for robust biological inference.

## Acknowledgements

We thank Lindsay K. Pino, Jian Wang, Ting Huang, and Joel R. Steele for thoughtful review this paper and analyses that contributed to poster presentation ahead of publication. We also thank the authors of the original datasets for making data available for re-use.

## Data Availability

The previously published MMCC dataset is available at ProteomeXchange (PXD042704), hosted via Panorama. The cancer dataset samples are available at ProteomeXchange (PXD060573), hosted via PRIDE. All result files from DIA-NN, scripts to generate figures, input and outputs to the `calculate-loq.py` script and the python script to generate peak sampled mzML files is available at ProteomeXchange PXD073415 hosted by MassIVE, MSV000100554. Examples of 100% concentration full, half, and quarter peak completeness files are provided alongside script. Help documentation embedded in the python peak sampling file will sufficiently enable re-creation of sampled files.

## Supplemental Figures

**Supplemental Figure S1.**
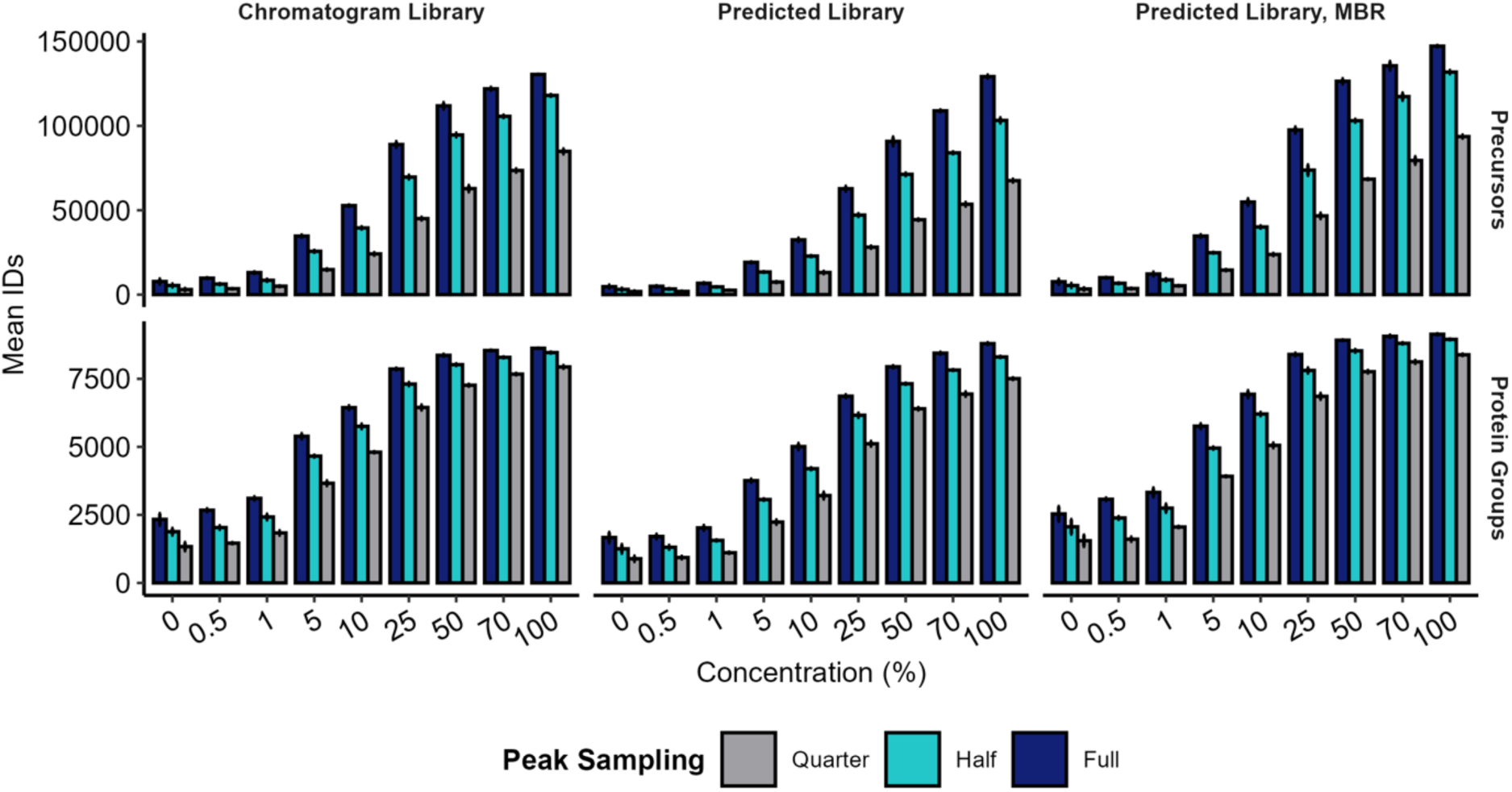
Mean precursor (top row) and protein-group (bottom row) identifications at 1% q-value across the MMCC concentration series for chromatogram library, predicted, and MBR workflows. Bar color indicates MS2 peak sampling completeness (full/half/quarter). This figure expands Figure 1A by showing all workflows side-by-side.

**Supplemental Figure S2.**
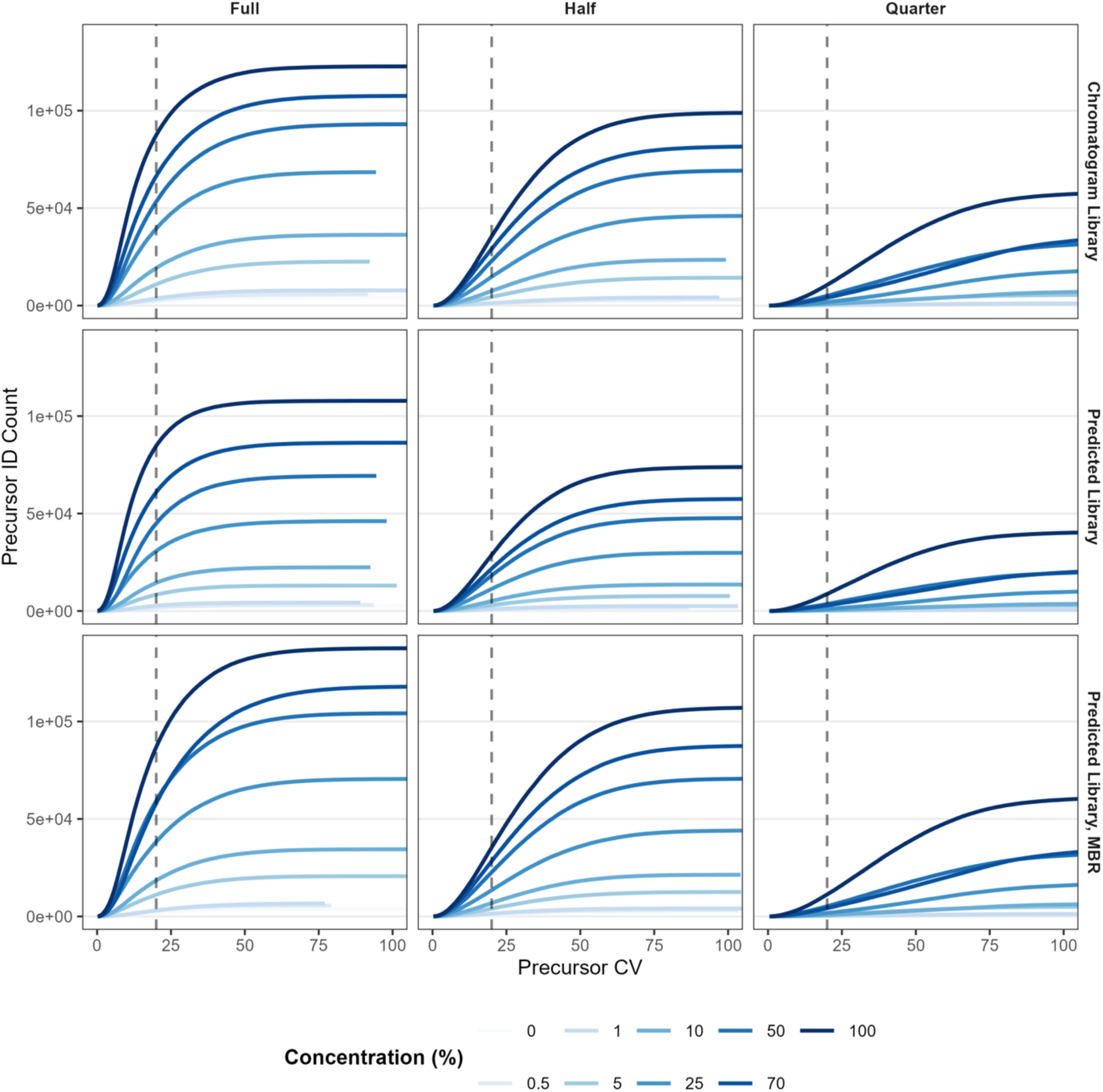
Precursor CV accumulation curves showing the number of identified precursors with CV ≤ threshold (x-axis) across MMCC concentrations (line color), faceted by peak sampling completeness (columns: full/half/quarter) and workflow (rows). The dashed vertical line marks CV = 20%, the threshold used for quantifiability comparisons.

**Supplemental Figure S3.**
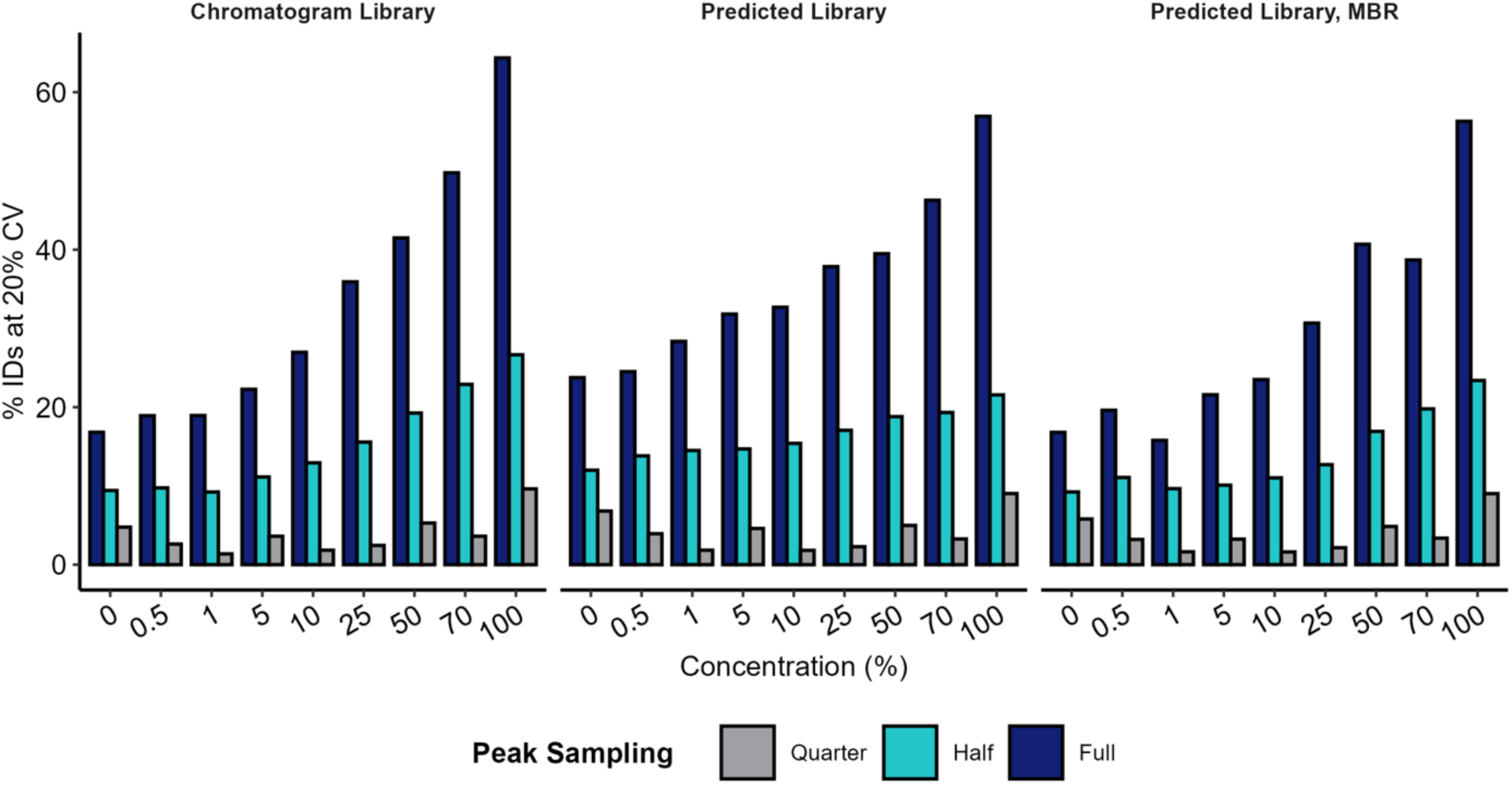
For each workflow, the percent of identified precursors with CV ≤ 20% is shown across MMCC concentrations. Bar color indicates peak sampling completeness (full/half/quarter). This figure expands Figure 1C and highlights the disproportionate loss of precision under reduced sampling, particularly at low concentrations.

**Supplemental Figure S4.**
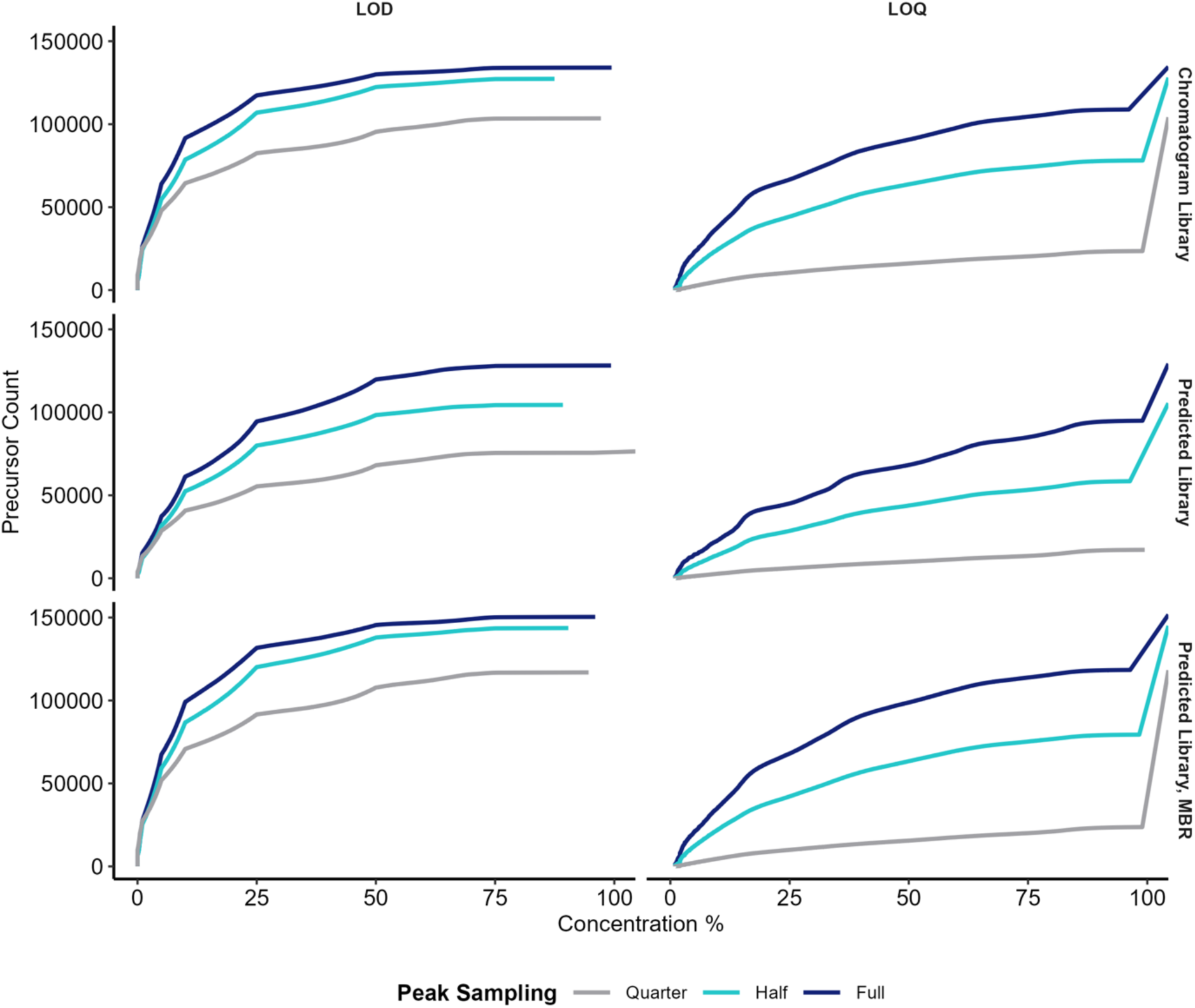
Accumulation plots showing the count of precursors whose LOD (left) or LOQ (right) is ≤ the concentration threshold on the x-axis, faceted by workflow (rows) with line series indicating sampling completeness (full/half/quarter). Values >100 correspond to precursors assigned “infinite” LOD/LOQ because the criterion is not met at any tested concentration; these are retained to preserve full distributions.

**Supplemental Figure S5.**
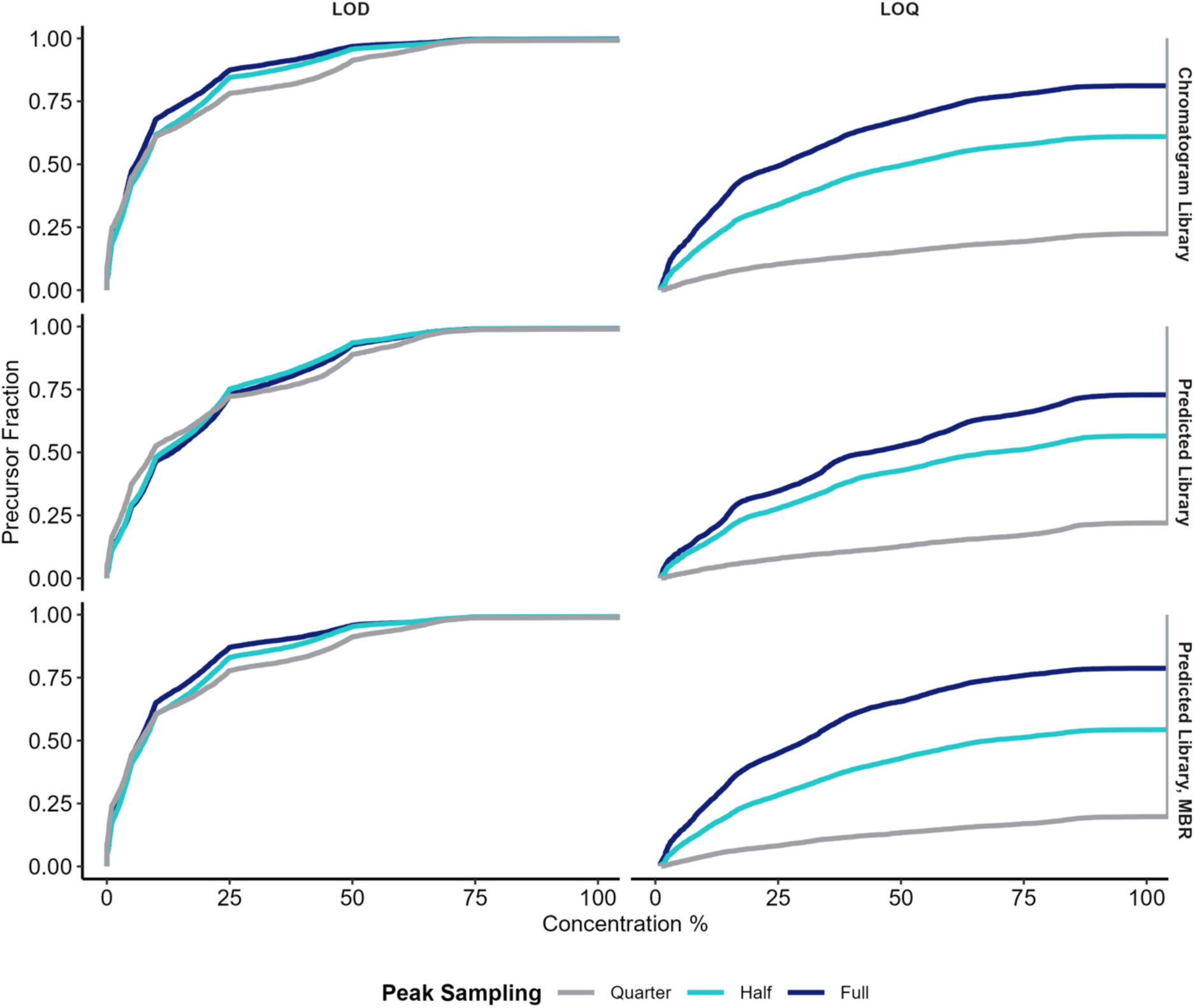
ECDFs of LOD and LOQ by workflow and peak sampling completeness. Empirical cumulative distribution functions (ECDFs) of precursor LOD (left) and LOQ (right), stratified by workflow (rows) and sampling completeness (full/half/quarter). Vertical behavior beyond 100% reflects precursors assigned infinite LOQ (detected but not meeting quantifiable linearity/precision criteria across the dilution series).

**Supplemental Figure S6.**
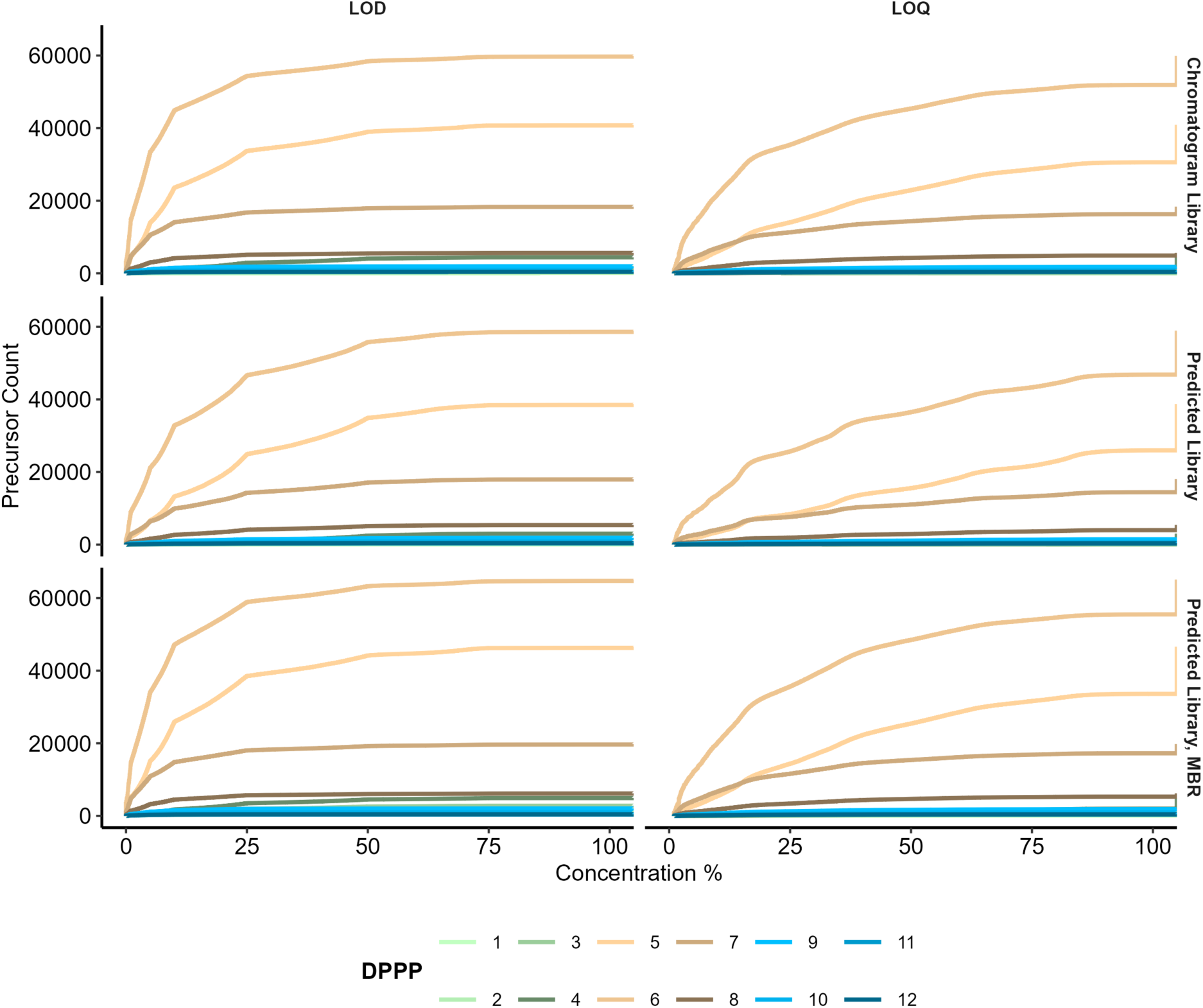
-Accumulation curves showing precursor counts meeting LOD (left) or LOQ (right) at or below the concentration threshold, with line series grouped by estimated DPPP and faceting by workflow (rows). This figure complements Figure 2C by emphasizing how quantifiability improves with sampling density, particularly between ∼3–6 DPPP for each workflow.

**Supplemental Figure S7.**
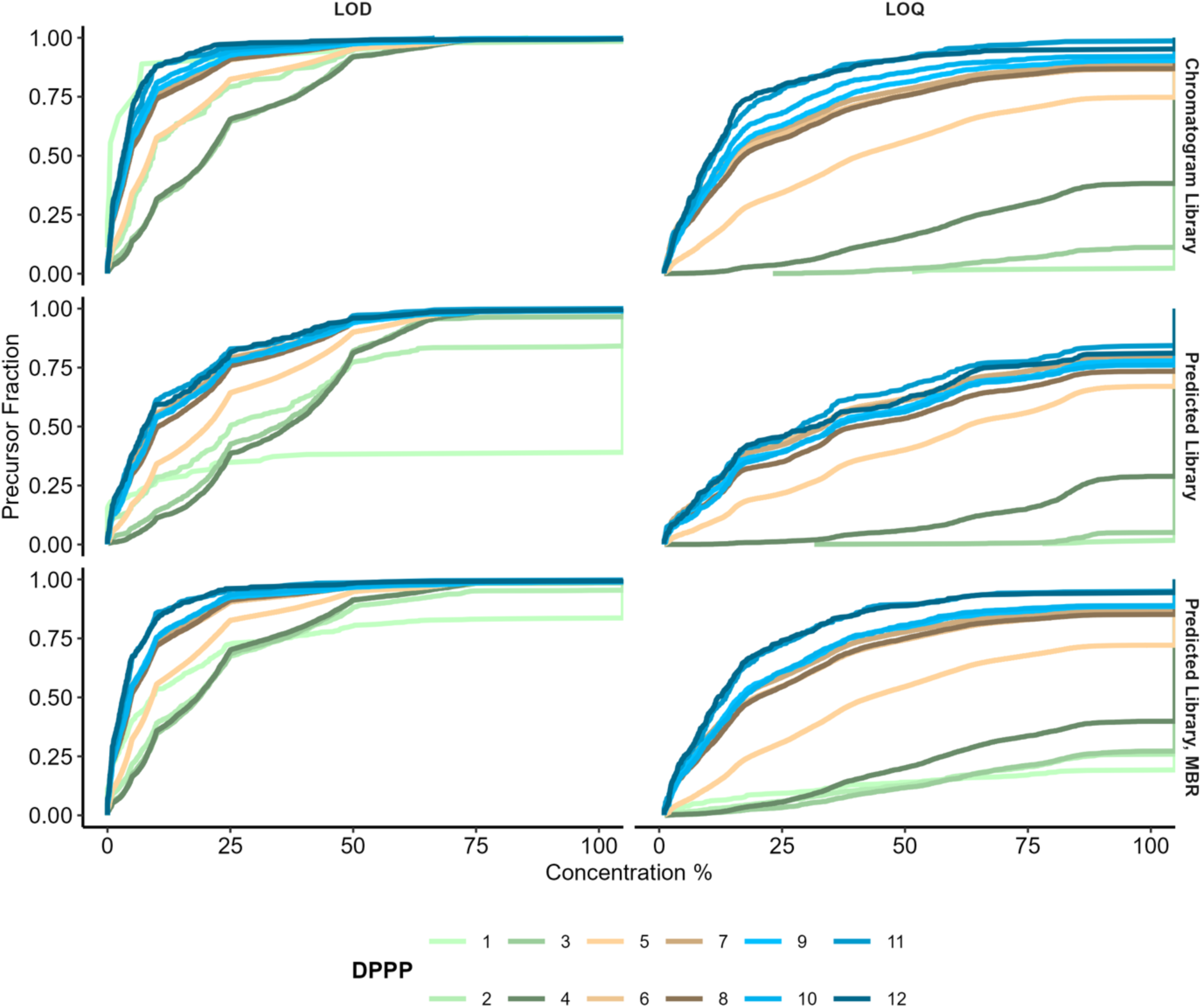
ECDFs of LOD (left) and LOQ (right), with line series grouped by estimated DPPP and faceted by workflow (rows). Lower curves indicate poorer sensitivity (higher LOD/LOQ). LOQ shows stronger DPPP-dependent separation than LOD across all workflows.

**Supplemental Figure S8.**
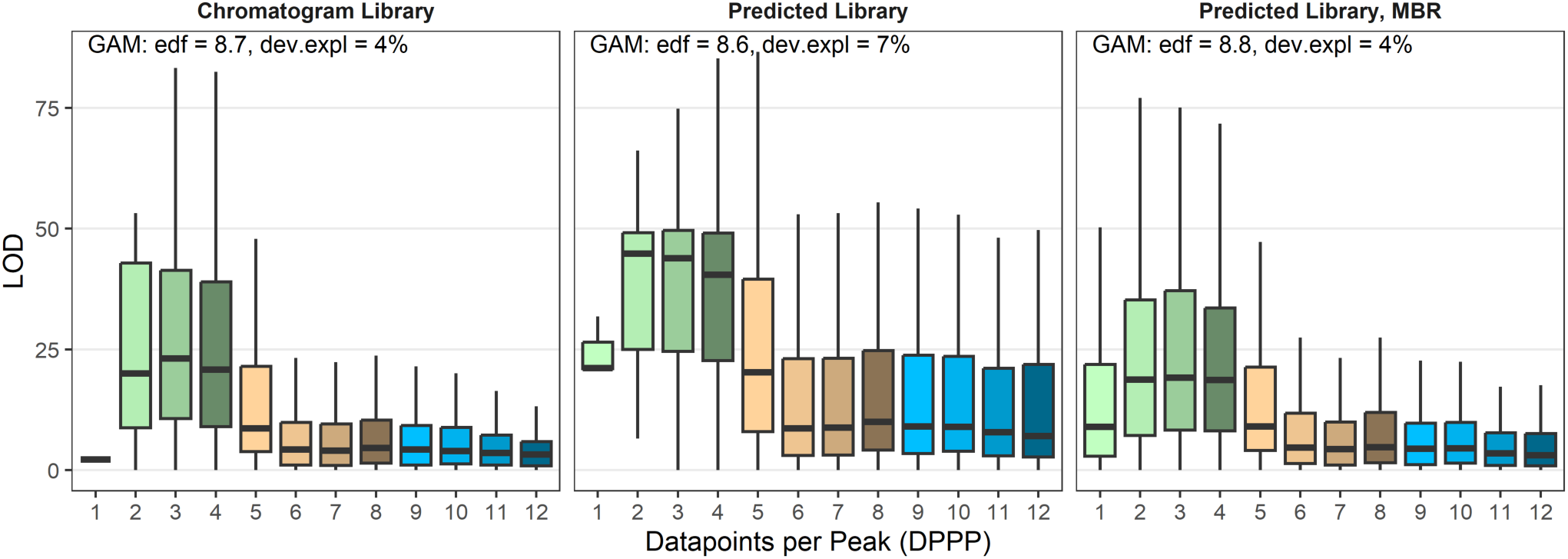
Boxplots of LOD as a function of discrete estimated DPPP for each workflow. GAM summaries (e[ective degrees of freedom and deviance explained) are shown for descriptive comparison, but deviance explained should be interpreted cautiously because DPPP is discrete and LOD values are bounded by the dilution series.

**Supplemental Figure S9.**
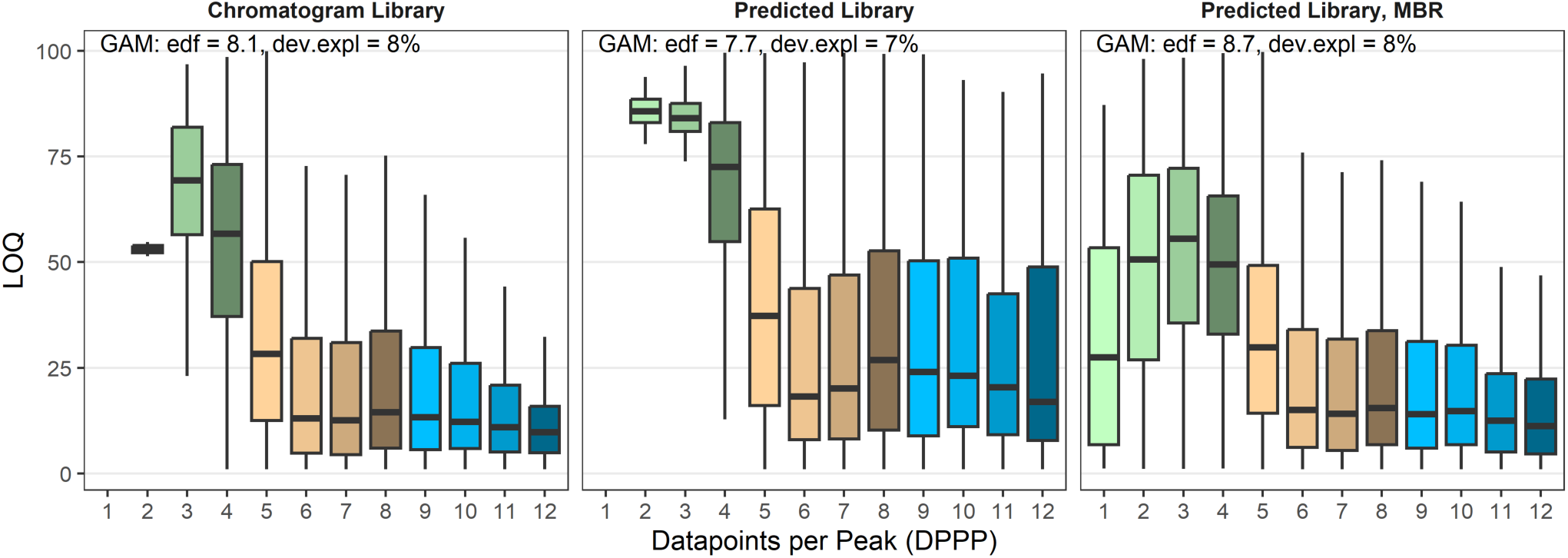
Boxplots of LOQ as a function of discrete estimated DPPP for each workflow. GAM summaries (e[ective degrees of freedom and deviance explained) are shown for descriptive comparison, but deviance explained should be interpreted cautiously because DPPP is discrete and LOQ values are bounded by the dilution series.

**Supplemental Figure S10.**
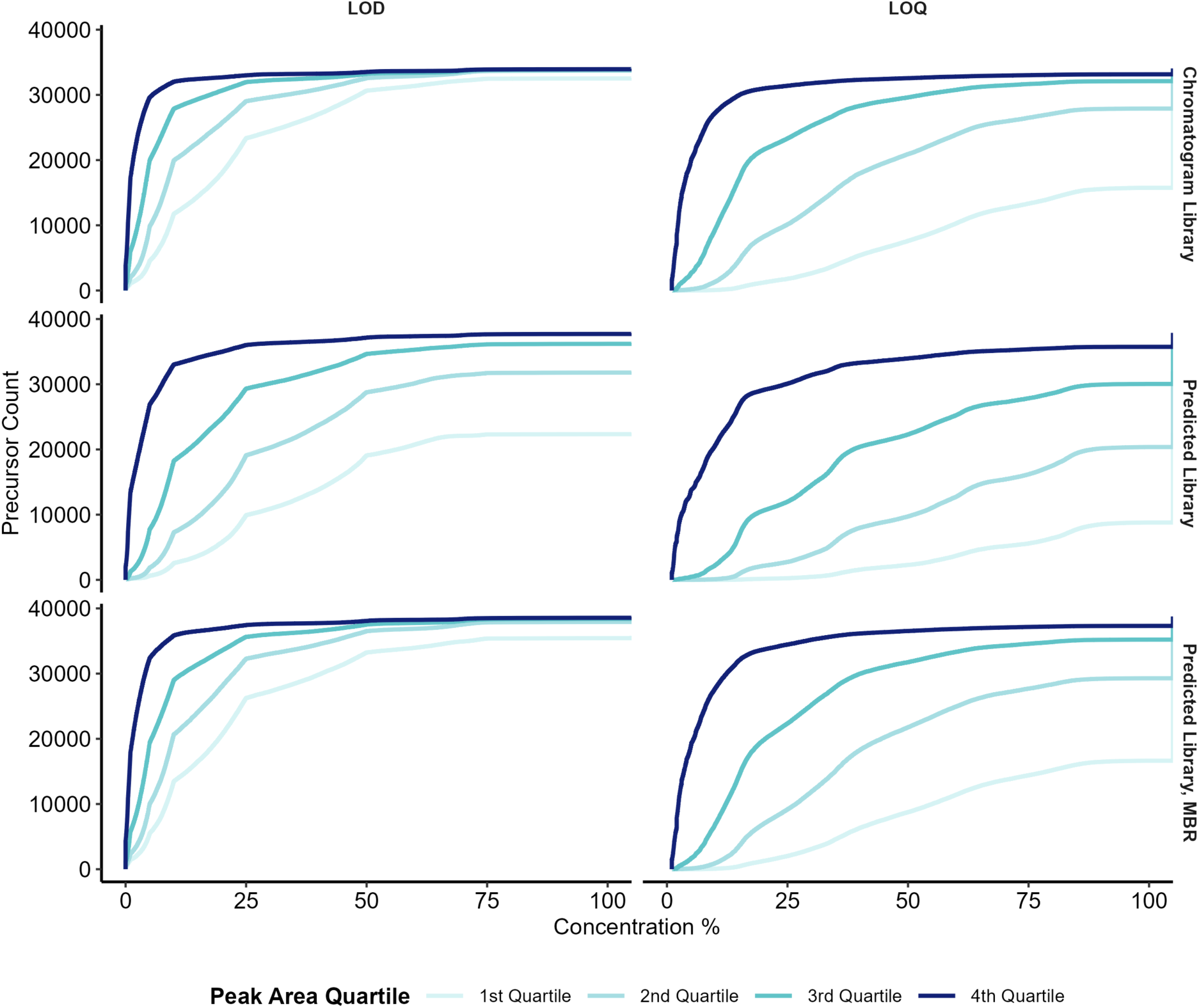
Accumulation curves of precursor counts meeting LOD (left) or LOQ (right) at or below the concentration threshold, stratified by peak area quartile at 100% concentration and faceted by workflow (rows). Because each quartile contains equal numbers of precursors, these curves compare count accumulation rather than fractional ECDF behavior.

**Supplemental Figure S11.**
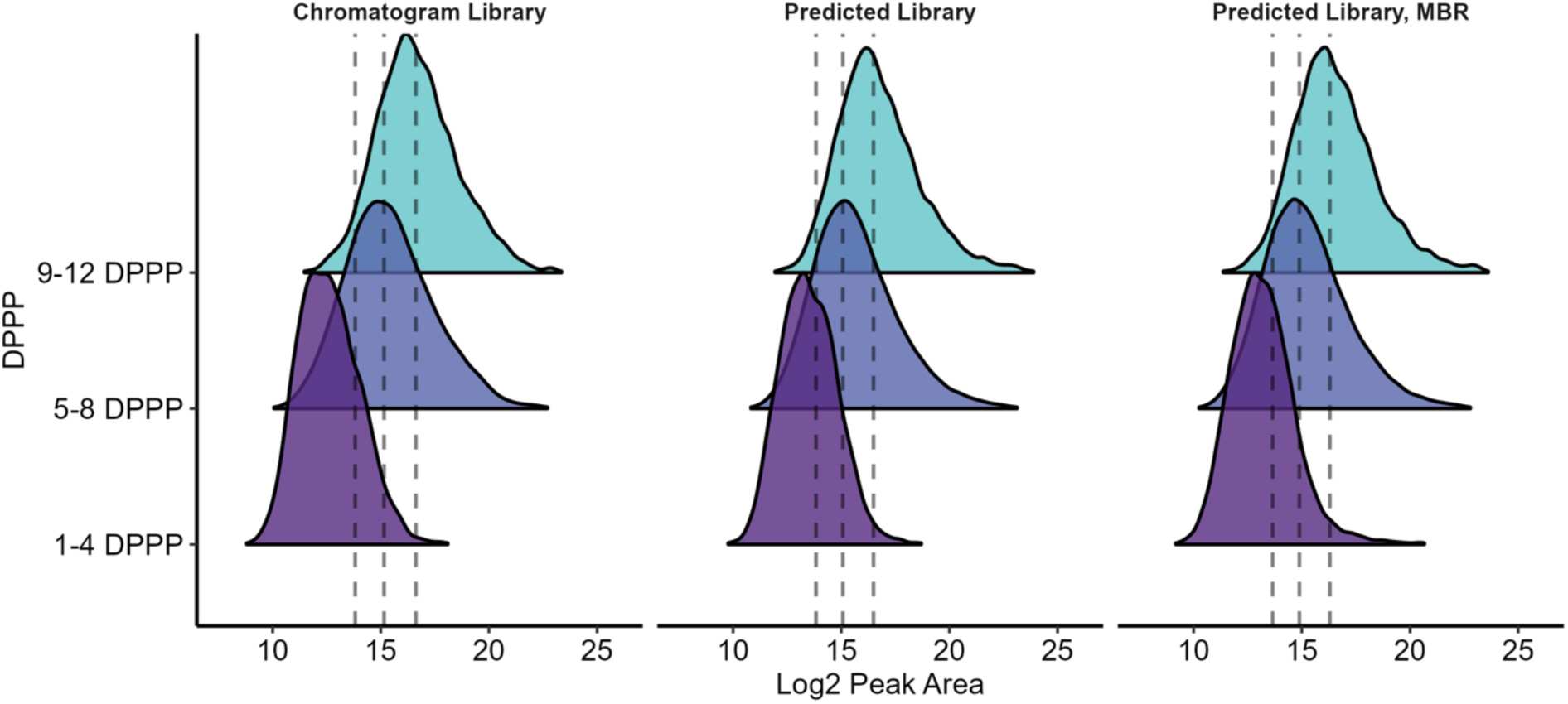
Ridgeline density plots of log2 precursor peak area (x-axis) for three grouped DPPP regimes (1–4, 5–8, 9–12), faceted by workflow. Vertical dashed lines denote the within-workflow peak area interquartile range (IQR). This visualization highlights that low-DPPP features are enriched at low peak area but can occur across the dynamic range.

**Supplemental Figure S12.**
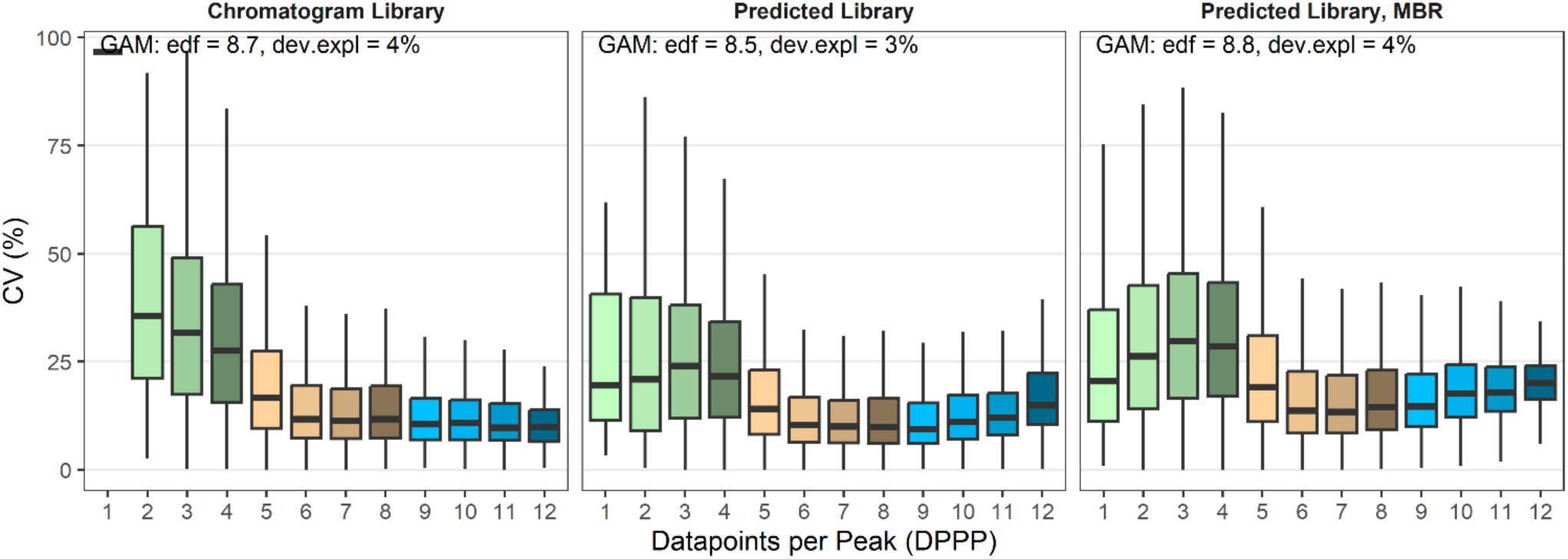
Boxplots showing precursor CV (%) versus estimated DPPP, faceted by workflow. Because DPPP is discrete/ordinal, smooth regression is not shown. GAM summaries are displayed for descriptive comparison of association strength.

**Supplemental Figure S13.**
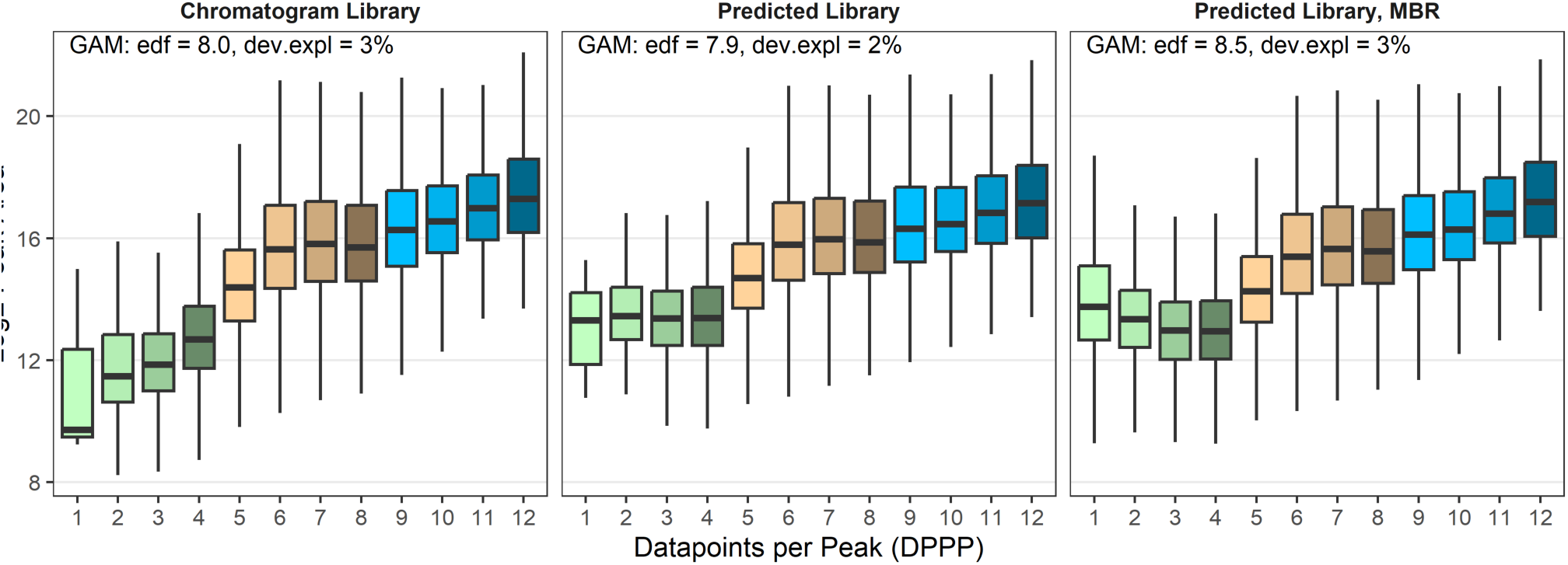
Boxplots of log_2_ precursor peak area at 100% concentration as a function of discrete DPPP, faceted by workflow. GAM summaries are shown for descriptive comparison; smooth curves are omitted because DPPP is discrete.

**Supplemental Figure S14.**
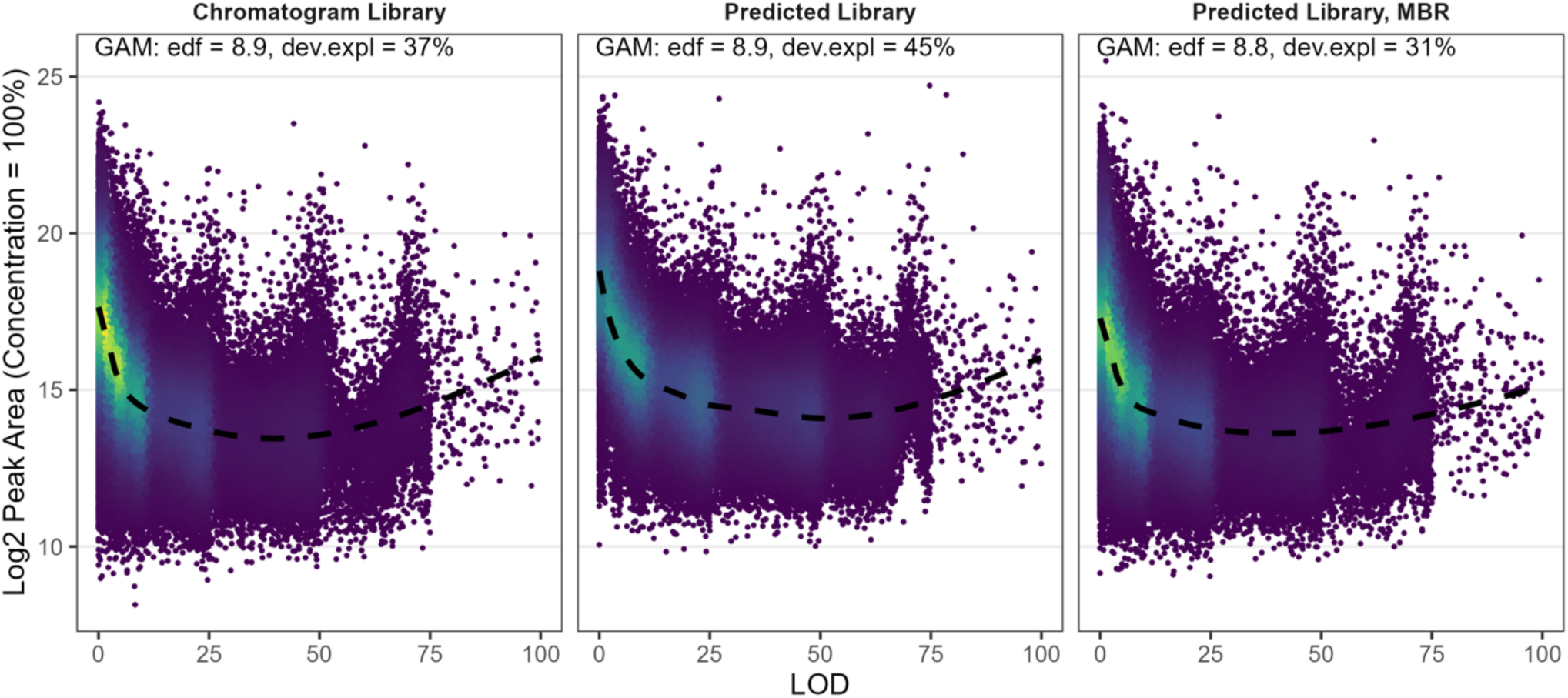
2D density plots of LOD (x-axis) versus log_2_ precursor peak area at 100% concentration (y-axis), faceted by workflow. Dashed curves show GAM smooth fits. This figure illustrates that detectability improves with peak area but saturates at higher signal levels.

**Supplemental Figure S15.**
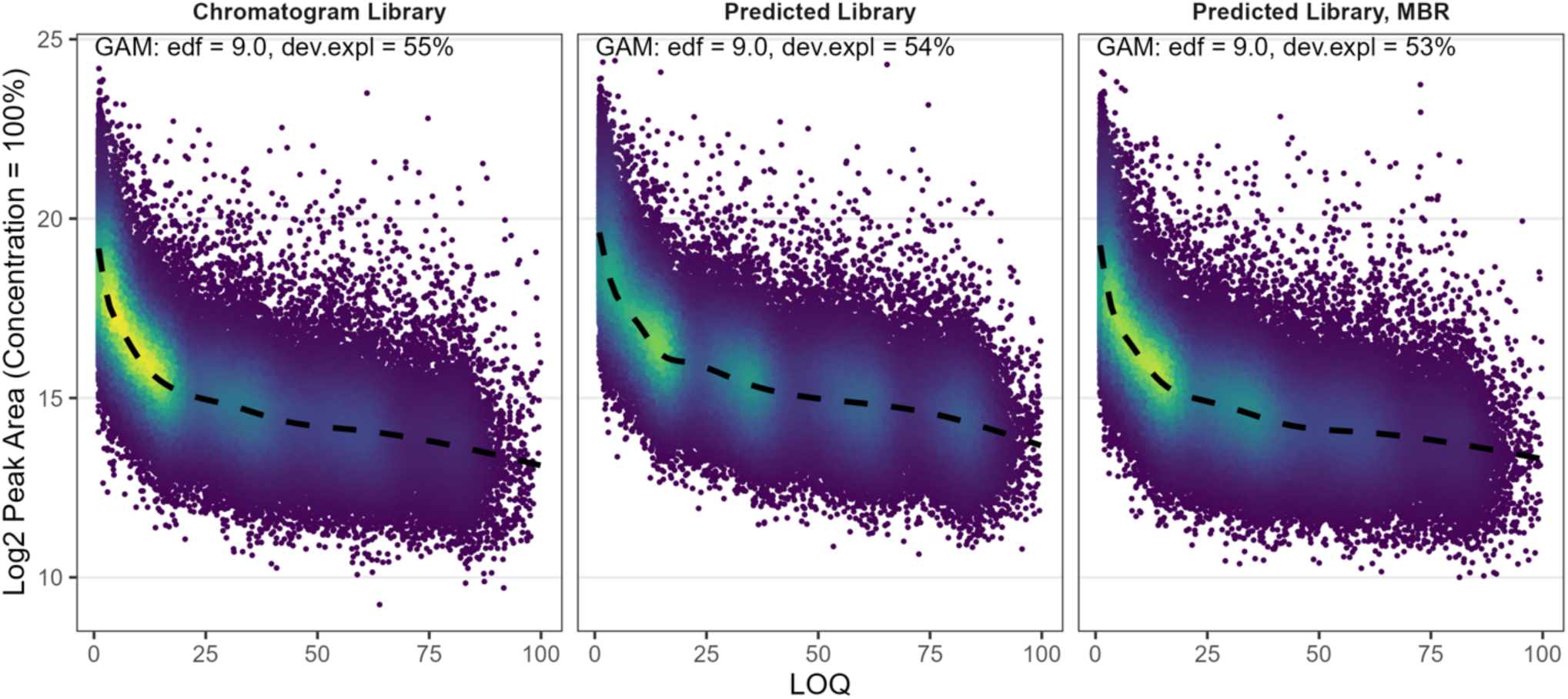
Density distribution of LOQ relative to log_2_ peak area at 100% concentration. Smoothed distribution drawn. The strongest nonlinear association observed in the study is between LOQ and peak area, supporting peak area as a primary driver of quantifiability.

**Supplemental Figure S16.**
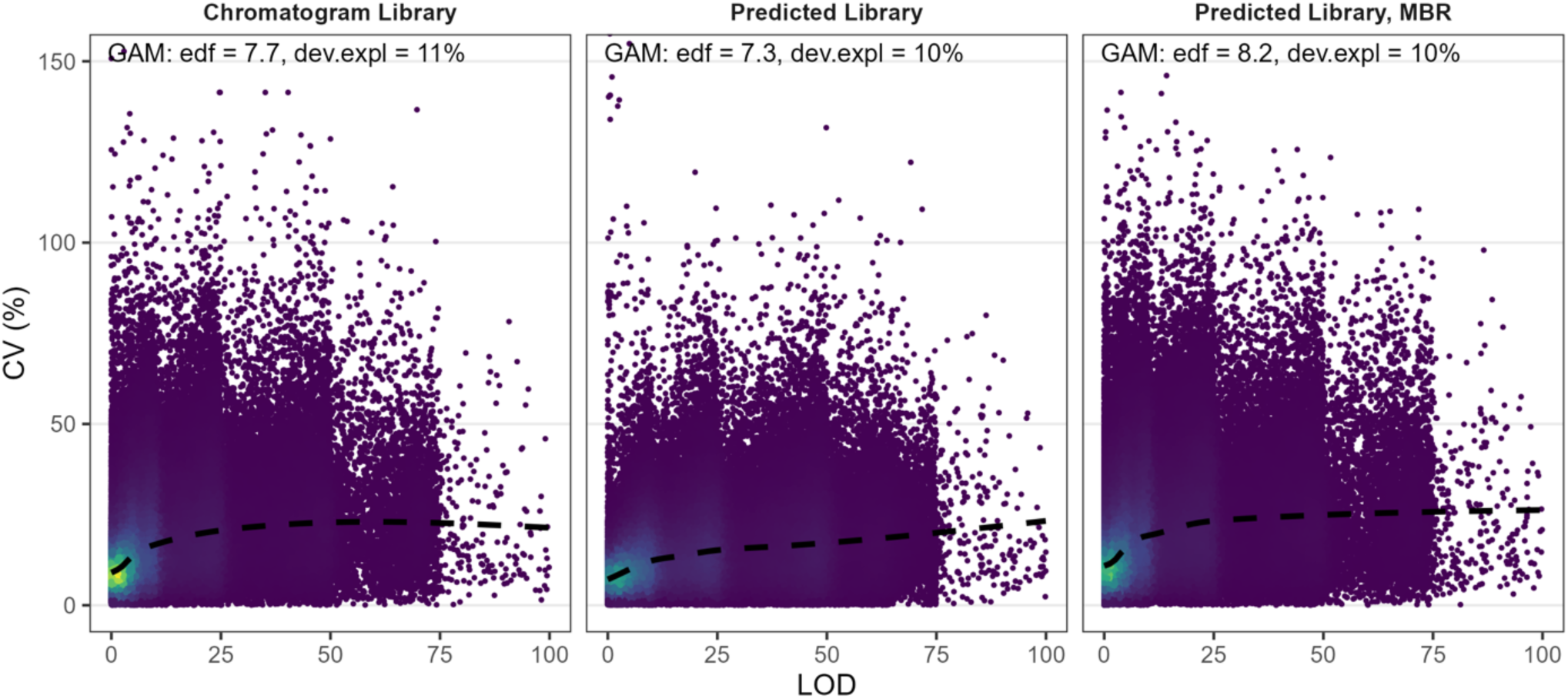
2D density plots of LOD (x-axis) versus precursor CV at 100% concentration (y-axis), faceted by workflow, with GAM smooth fits shown as dashed lines. This visualization supports that LOD is primarily a detectability metric and is less directly governed by precision than LOQ.

**Supplemental Figure S17.**
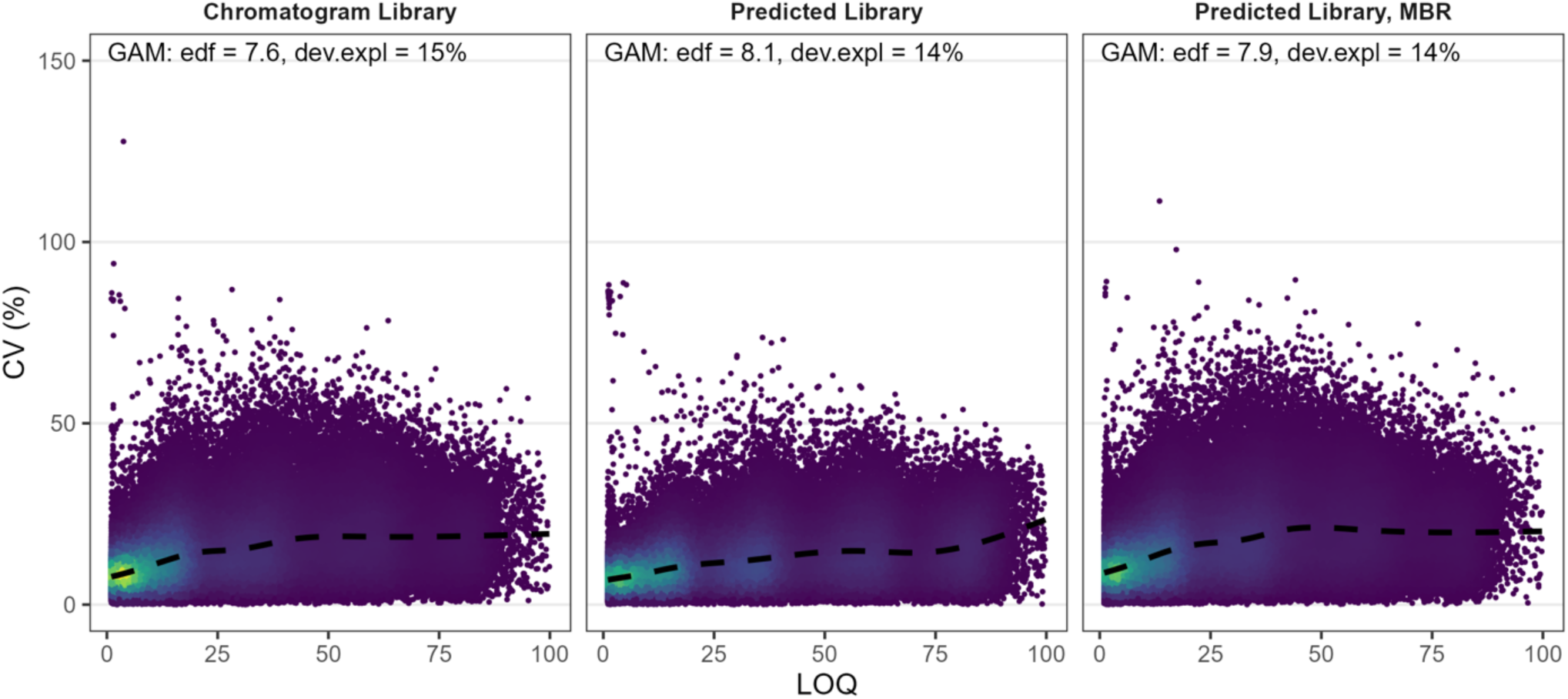
2D density plots of LOQ (x-axis) versus precursor CV at 100% concentration (y-axis), faceted by workflow, with GAM smooth fits shown as dashed lines. Higher CV is associated with worse LOQ, reflecting the LOQ requirement for both linearity and CV ≤ 20% under resampling.

**Supplemental Figure S18.**
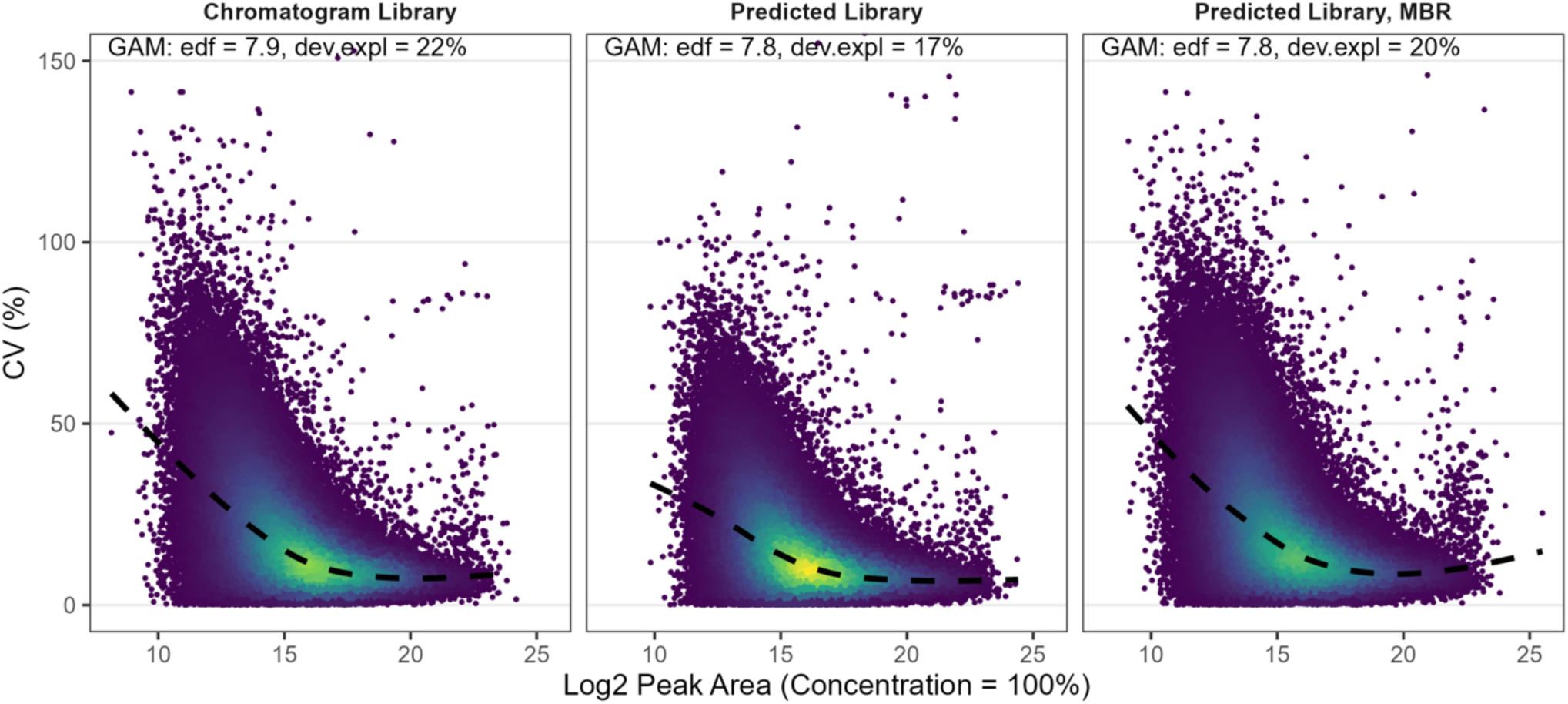
2D density plots of precursor CV (%) (y-axis) versus log_2_ precursor peak area at 100% concentration (x-axis), faceted by workflow, with GAM smooth fits shown as dashed lines. This figure contextualizes that higher-abundance features tend to exhibit improved precision, linking abundance context to quantifiability.

**Supplemental Figure S19.**
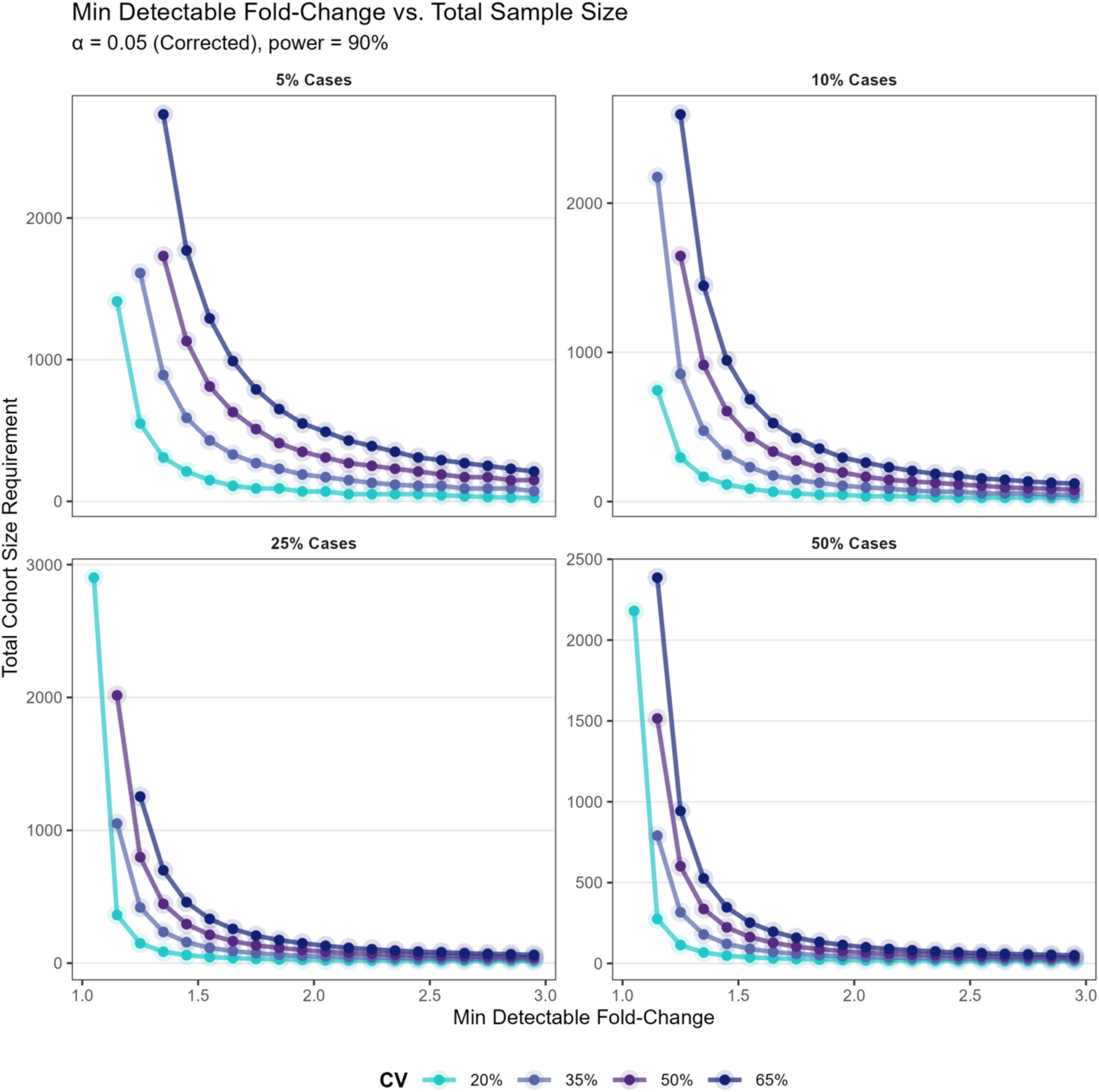
Minimum detectable fold-change required to achieve 90% power (y-axis) as a function of total sample size (x-axis), simulated for cohorts with di[erent case fractions (panels) while measuring 3,000 protein groups. Line series represent assumed technical CV (20–65%). Increased CV substantially increases the fold-change required for adequate power, particularly in unbalanced designs where e[ective sample size is limited.

**Supplemental Figure S20.**
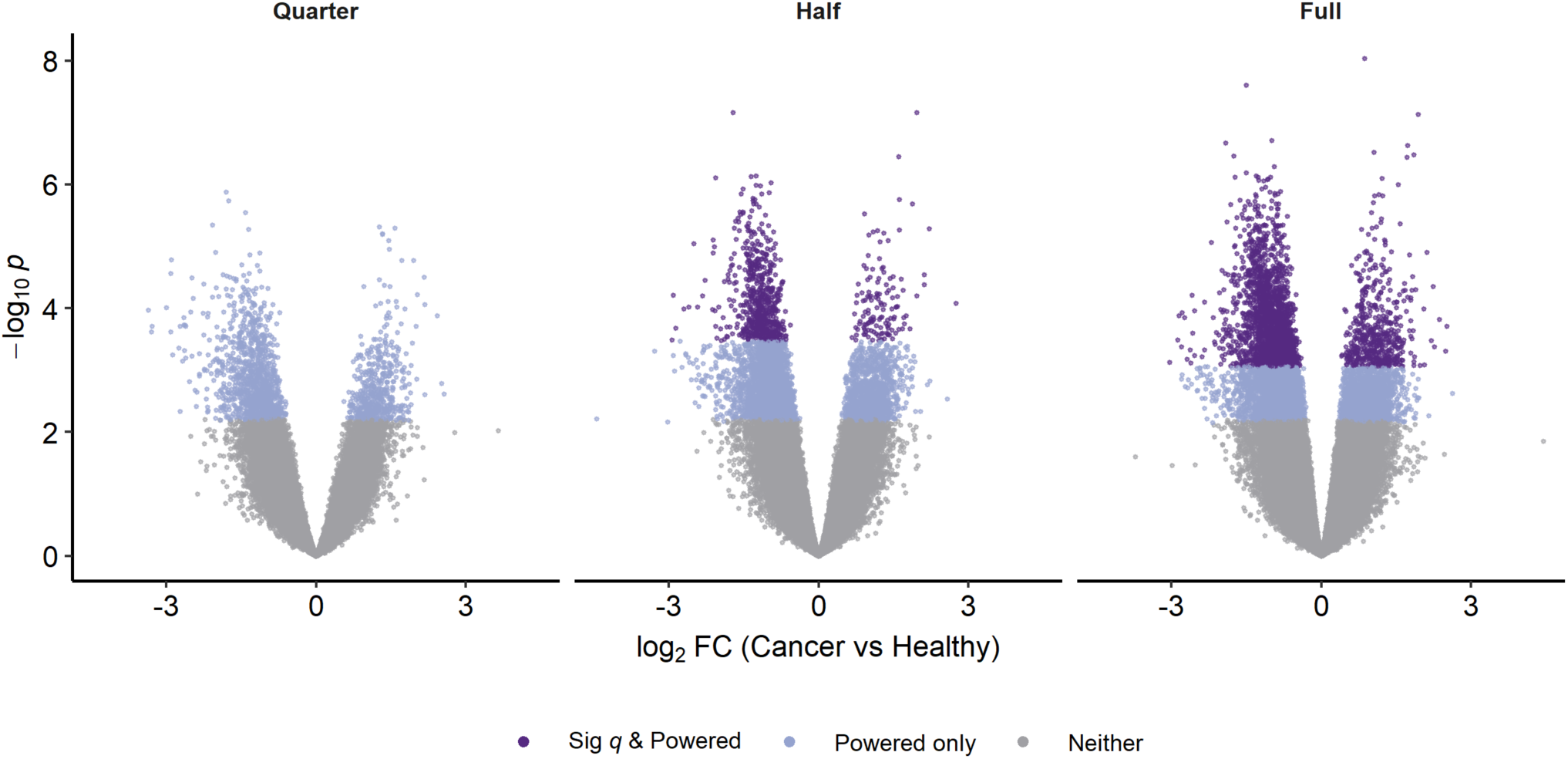
Volcano plot accompanying Figure 5.

**Supplemental Figure S21.**
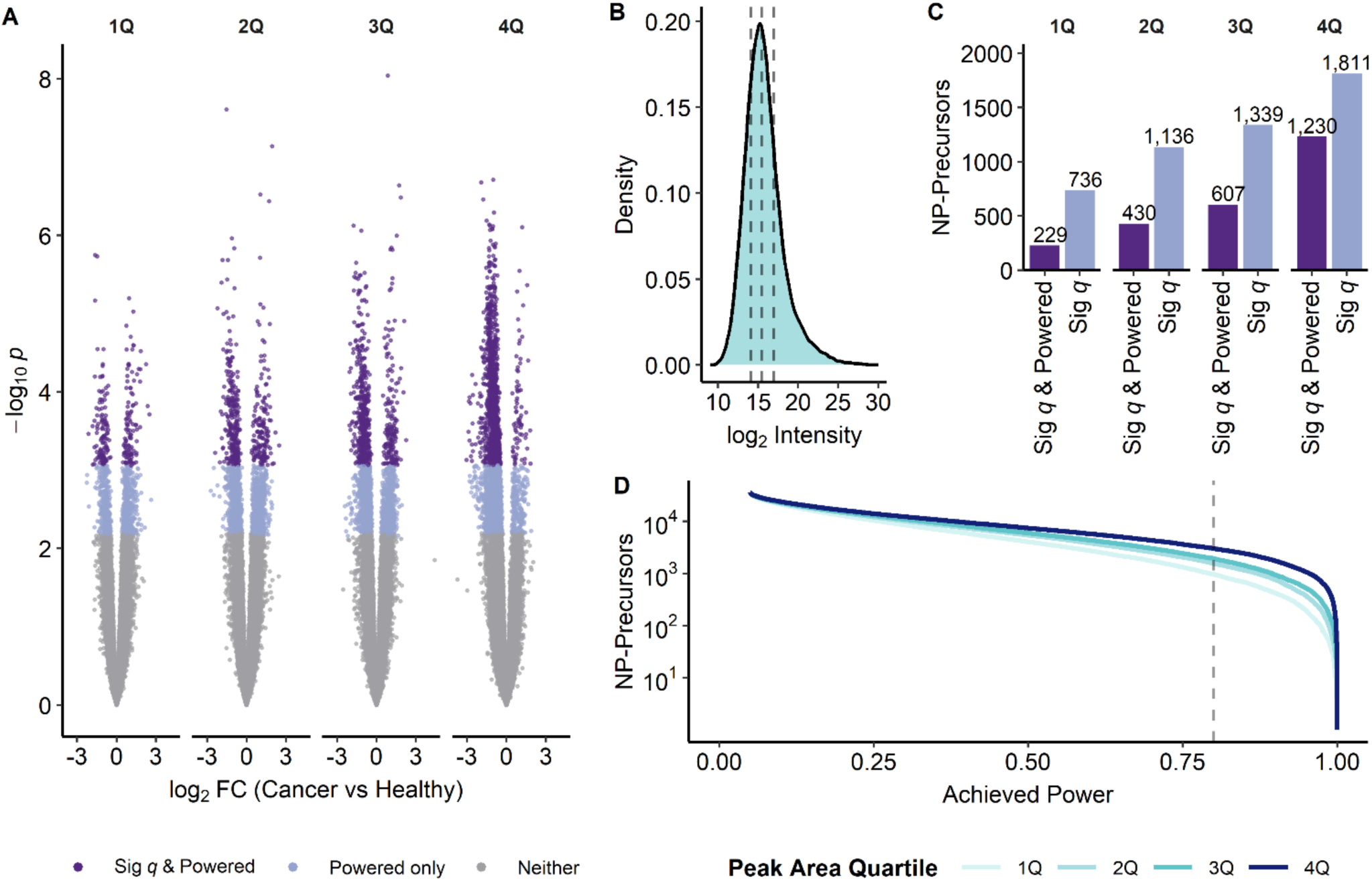
(A) Volcano plots of cancer vs healthy differential abundance results stratified by peak area quartile using post-filtering precursor peak area. Points are colored by BH significance and achieved power classification. (B) Density distribution of the peak area metric used for quartile assignment with dashed lines marking quartile boundaries. (C) Counts of BH-significant NP-precursors and BH-significant + powered NP-precursors by quartile; each quartile contains equal precursor counts enabling direct comparison. (D) Power accumulation curves by quartile with the 0.80 power threshold indicated.

